# Functional recovery of vitrified and nanowarmed human-scale pig kidneys

**DOI:** 10.64898/2026.10.07.756990

**Authors:** Zonghu Han, Bat-Erdene Namsrai, Anudari Chimedtseren, Anthony A. D. Chen, Benjamin J. Fisher, Diane Tobolt, Michael G. Megaly, Onyinyechukwu Justina Oziri, Minhan Jiang, Joseph Sushil Rao, Andrew B. Adams, Michael L. Etheridge, John C. Bischof, Erik B. Finger

## Abstract

Organ transplantation is critically constrained by the short preservation window for donor organs, limiting allocation and contributing to organ discard and waitlist mortality. Cryogenic organ banking could extend preservation from hours to months or years, but ice formation, thermal stress and cryoprotectant toxicity become increasingly difficult to control as organ size increases. Vitrification and nanowarming have enabled long-term preservation and transplantation of rodent kidneys but have not been demonstrated at the scale of clinically relevant organs. Here we show that vitrification and electromagnetic nanowarming enable long-term cryogenic storage and functional recovery of human-scale porcine kidneys. Kidneys 100-fold larger than previously nanowarmed kidneys were successfully recovered after storage at −150 °C for up to one year. A new cryoprotectant formulation and multithermic perfusion strategy enabled ice-free vitrification at cooling rates achievable in large kidneys, while controlled cooling with annealing and staged volumetric nanowarming limited thermal stress and enabled rapid rewarming without cracking. Rewarmed kidneys demonstrated vascular, metabolic and excretory function during normothermic perfusion, with limited histological injury, and reperfused homogeneously and produced urine immediately after transplantation. Functional recovery approached static cold storage outcomes across several measures and was similar to that of kidneys exposed to cryoprotectant alone, suggesting minimal additional functional injury from vitrification, cryogenic storage and nanowarming. These findings overcome major scale and storage-duration barriers to organ cryopreservation and establish a path toward organ banking on timescales far beyond those of conventional preservation.

## Introduction

The inability to preserve donor organs beyond hours remains a fundamental limitation in transplantation. Static cold storage (SCS) slows metabolic deterioration but cannot arrest it indefinitely ^1^, constraining organ allocation and transport as cold ischemia time accumulates. Cryogenic organ banking could extend preservation from hours to months or years ^2^, enabling more flexible donor-recipient matching and allocation, while providing time for organ assessment, recipient preparation and therapeutic intervention prior to transplantation.

Efforts to cryopreserve whole organs date back more than half a century, but conventional freezing approaches were limited by ice formation, osmotic injury and tissue disruption ^3^. Vitrification, the conversion of a liquid into an ice-free glassy state, offers an alternative by suppressing ice formation during cooling ^3,4^. This approach ultimately enabled vitrification, rewarming and transplantation of a rabbit kidney, but also highlighted persistent limitations related to cryoprotective agent (CPA) delivery, toxicity and thermal constraints ^5^. In particular, the critical warming rate (CWR) required to prevent devitrification during rewarming can greatly exceed the critical cooling rate (CCR) required to avoid ice during vitrification ^6^. As specimen size increases, surface warming becomes increasingly slow and non-uniform, leading to ice formation within the organ core while generating thermal gradients that can cause cracking ^7^. Increasing CPA concentrations can improve resistance to ice formation but increase concentration- and temperature-dependent toxicity ^8,9^. Whole-organ vitrification therefore requires simultaneous control of ice formation, thermal stress and CPA toxicity.

Nanowarming was developed to overcome the limitations of surface-based rewarming ^10,11^. Silica-coated iron oxide nanoparticles (sIONPs) distributed through the vasculature before vitrification generate heat when exposed to an alternating electromagnetic field, enabling rapid volumetric rewarming throughout the organ. We extended nanowarming to explanted rat kidneys, enabling rapid rewarming with reduced physical and biological injury compared with convective warming ^12^. Further refining this approach, we subsequently demonstrated vitrification, long-term cryogenic storage (up to 100 days), nanowarming and life-sustaining transplantation of rat kidneys (∼1 g) ^13^. More recently, physical vitrification and nanowarming were demonstrated at liter-scale volumes approaching those of human organs ^14^. Whether an integrated vitrification and nanowarming process could preserve biological function in a human-scale organ, however, remained unknown.

Scaling from rat to human-scale porcine kidneys introduces additional constraints during both cooling and CPA delivery. As organ size increases, conductive heat transfer limits the cooling rate achievable within the organ core ^14^. Under these conditions, the CPA formulation used for rat kidney vitrification (VMP) does not provide sufficient protection against ice formation, particularly in incompletely equilibrated regions of the kidney ^15^. Larger organs also require longer CPA loading and unloading, increasing exposure to potentially toxic CPA concentrations, while heterogeneous renal perfusion can limit equilibration in regions such as the medulla ^5,16^. Scaling vitrification therefore requires coordinated optimization of CPA composition and delivery, cooling, nanoparticle distribution and nanowarming.

Here, we developed an integrated approach to vitrification and nanowarming of human-scale porcine kidneys. We combined a new CPA formulation designed for the slower cooling rates of large organs with multithermic CPA delivery, balanced vascular sIONP loading, controlled cooling and volumetric nanowarming. We assessed physical performance; vascular, metabolic and excretory function; and structural integrity during normothermic machine perfusion. Comparisons with fresh, 24-h SCS and CPA-only kidneys distinguished injury associated with CPA exposure and delivery from additional injury associated with vitrification, cryogenic storage and nanowarming. Finally, we performed acute allotransplantation to assess reperfusion and immediate function in vivo. Together, these studies test whether long-term cryogenic organ banking can be extended from small-animal proof of concept to organs at a scale relevant to human transplantation.

## Results

### Scaling vitrification and nanowarming to human-scale kidneys

We extended vitrification and nanowarming (VNW) from ∼1-g rat kidneys to human-scale porcine kidneys (∼100 g, range 84–132 g, Extended Data Table 1), representing an approximately 100-fold increase in organ mass (Fig. 1a). The VNW process comprised cryoprotective agent (CPA) loading, vascular delivery of silica-coated iron oxide nanoparticles (sIONPs), cooling to a vitrified state, cryogenic storage, volumetric nanowarming, CPA and sIONP unloading and subsequent functional assessment or transplantation (Fig. 1b).

**Fig. 1.**
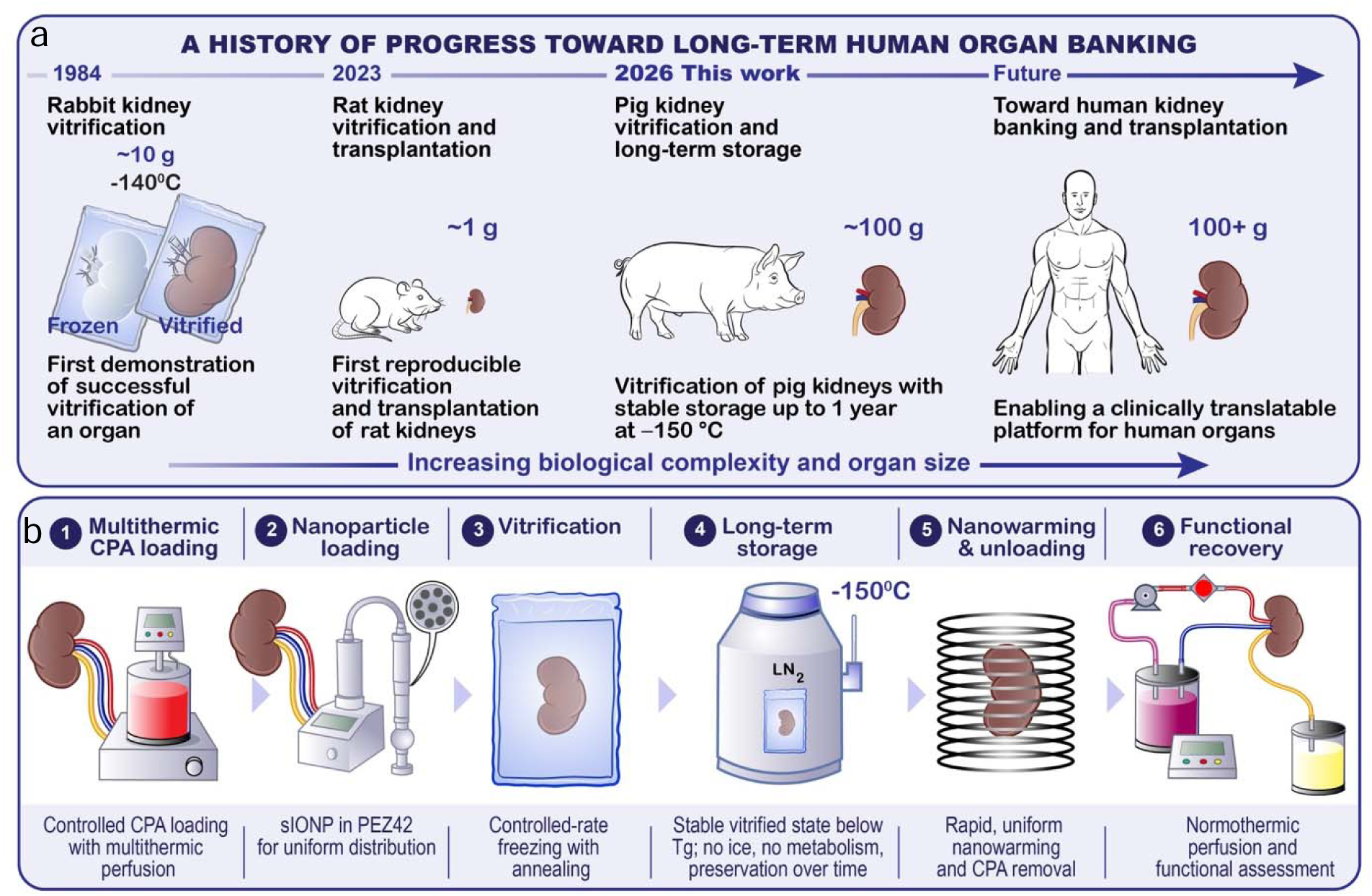
Overview of vitrification and nanowarming of porcine kidneys. **a,** Historical progression of organ cryopreservation from the development of vitrification in the 1980s ^1^, through successful vitrification, nanowarming and transplantation of rat kidneys ^2^, to the present study scaling vitrification and nanowarming to porcine kidneys. **b,** Experimental workflow for whole-kidney vitrification and nanowarming (VNW), including cryoprotective agent (CPA) loading, silica-coated iron oxide nanoparticle (sIONP) loading, vitrification, cryogenic storage, nanowarming, CPA unloading and subsequent assessment by normothermic machine perfusion (NMP) and transplantation. PEZ42, vitrification solution used in the present study; Tg, glass transition temperature.

Scaling VNW to porcine kidneys required substantial development beyond protocols successful in the rat kidney. Reduced cooling rates within the organ core required a CPA with greater resistance to ice formation, while longer CPA loading and unloading increased the potential for toxicity. We therefore developed a higher-stability CPA formulation (PEZ42), combined it with multithermic perfusion to limit CPA toxicity and adapted cooling and nanowarming for human-scale kidneys. We evaluated the resulting protocol through physical characterization, normothermic machine perfusion, histological assessment and acute allotransplantation.

### Development of a CPA for human-sized kidneys

Thermal conduction constraints within 100-g porcine kidneys restrict core cooling rates to ∼2 °C/min ^14^. The VMP formulation previously validated in rat kidneys has a critical cooling rate (CCR) of ∼5 °C/min and therefore cannot reliably vitrify under these large-organ cooling conditions. This limitation is compounded by heterogeneous CPA distribution within the kidney, particularly in the poorly perfused renal medulla. Prior studies of rabbit kidneys demonstrated that medullary CPA concentrations may reach only ∼92% of the arterial concentration ^5^, consistent with our modeling of CPA distribution in rat kidneys ^15,16^. At 92% concentration, the CCR of VMP increases to ∼15 °C/min.

Avoiding even small amounts of medullary ice is important for renal recovery. In vitrified rabbit kidneys, Fahy and colleagues estimated that formation of ∼1–2% ice in the inner medulla during rewarming was associated with substantial medullary injury, while subsequent studies showed that >2% inner-medullary ice caused selective destruction of the medullary vasculature and failure of medullary perfusion after transplantation ^5,17^. These observations establish a biological upper limit for tolerable ice formation but do not define the more stringent physical criterion conventionally used to characterize successful vitrification. For CCR determination by differential scanning calorimetry (DSC), successful vitrification is operationally defined as limiting ice formation to <0.2% ^3^. We therefore adopted <0.2% ice as our design criterion and required candidate formulations to remain below this threshold when cooled at 2 °C/min at 92% of full concentration.

We sought to achieve this physical stability while maintaining sufficient biological compatibility for the prolonged CPA loading required in a large organ. M22, developed by Fahy and colleagues for organ vitrification, provided an important benchmark, with exceptionally low critical cooling and warming rates (CCR, 0.1 °C/min; CWR, 0.4 °C/min) at full strength (64.8% w/v) ^3^. However, M22 showed more chemical toxicity than VMP in our initial kidney studies, a critical limitation given the prolonged loading times required for large organs. We therefore developed a new CPA formulation intended for human-scale kidneys. Using M22 and VMP as performance benchmarks, and after several generations of optimization including mass transport modeling, empiric testing and validation in rat kidneys, we identified PEZ42. Relative to VMP, PEZ42 replaces formamide, a relatively weak glass former, with the stronger glass former propylene glycol and increases ethylene glycol by 4% w/v and the non-penetrating ice blocker Z-1000 by 2% w/v. DSC showed that both 100% and 92% PEZ42 limited ice formation to <0.2% during cooling at 1 °C/min, meeting the predefined vitrification criterion even at the concentration expected in the renal medulla. During rewarming, the CWR decreased with increasing PEZ42 concentration, from 2.3 °C/min at 92% to 1.3 °C/min at 96% and <1 °C/min at 100% (Supplementary Fig. 1). These preliminary data supported progression of PEZ42 to large-organ evaluation.

### Optimization of multithermic CPA and nanoparticle loading

To mitigate chemical toxicity from high solute concentrations (∼7.95 M), we implemented a dynamic multithermic machine perfusion approach. In initial trials, perfusion with full-strength CPA at 4 °C caused severe toxicity and functional failure during NMP. Leveraging observations that subzero temperatures (≤ −15 °C) suppress chemical toxicity ^5^, we designed a temperature-modulated protocol.

CPA loading was initiated at 4 °C with low CPA concentrations in LM6 carrier solution (Methods) to maintain low vascular resistance, with CPA concentration gradually increased to limit osmotic volume excursions. As the kidneys transitioned to higher CPA concentrations, first to an intermediate CPA (PMP) and then to PEZ42, the perfusate temperature was progressively lowered to below −15 °C to suppress CPA toxicity (Fig. 2a). Perfusion pressure was increased from 40 mmHg at 4 °C to 60 mmHg during PMP loading and to 80 mmHg during PEZ42 loading to compensate for increasing fluid viscosity and maintain tissue perfusion. Exposure to full-strength PEZ42 was minimized by monitoring ureteral effluent CPA concentration by refractometry and terminating loading once the effluent reached ≥97.5% of full strength (Fig. 2a–c; Supplementary Fig. 2).

**Fig. 2.**
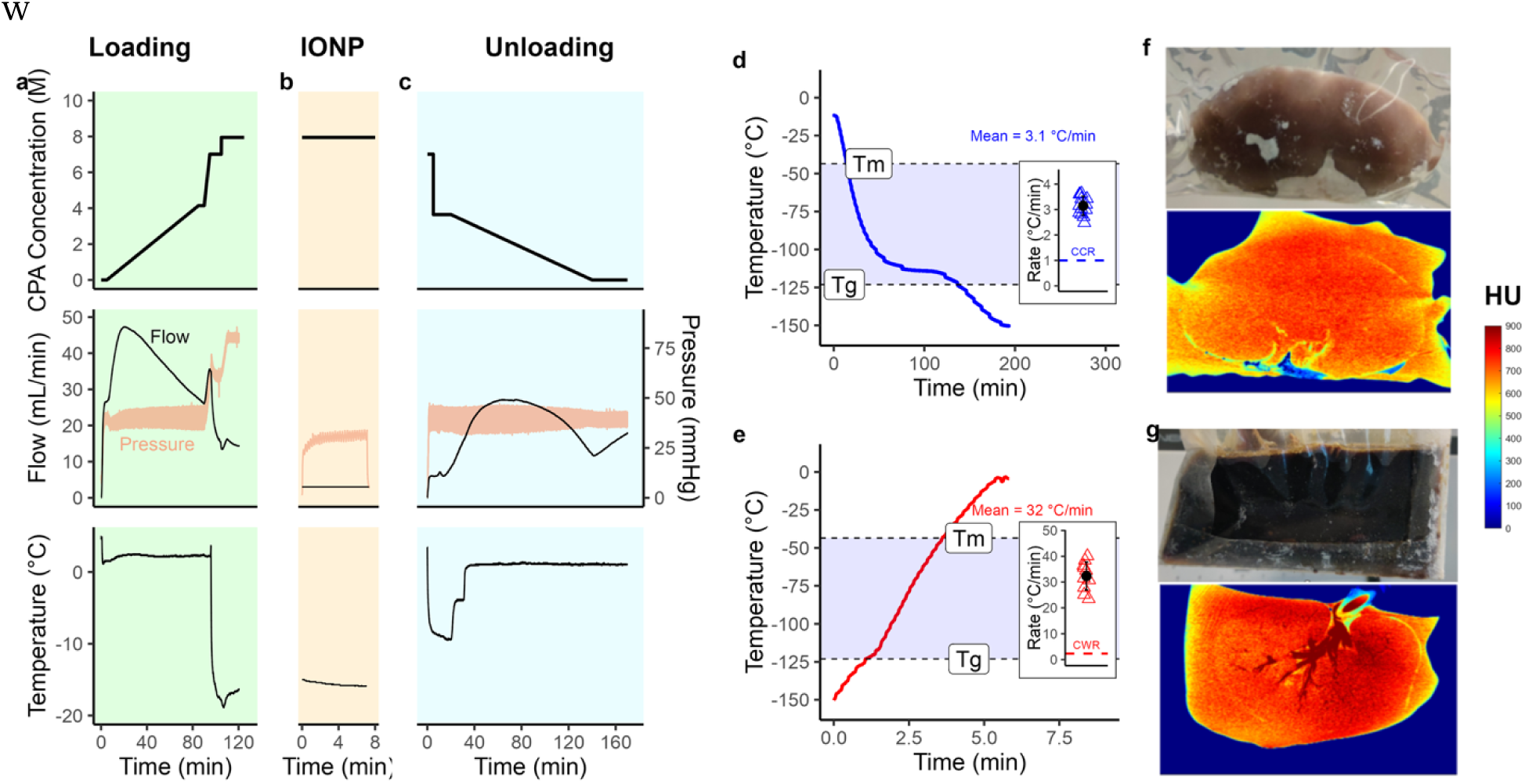
Whole-kidney vitrification and nanowarming achieve ice-free cryopreservation. Whole-kidney cryoprotective agent (CPA) loading, vitrification and nanowarming achieved the physical requirements for ice-free cooling and rewarming. Panels a–c, f and g show representative kidneys and illustrate the protocol and physical outcomes; insets in d and e show all kidneys used to characterize cooling and nanowarming rates. **a,** CPA loading protocol showing perfusate CPA concentration (top), flow and pressure (middle) and kidney temperature (bottom) during the loading procedure. **b,** Silica-coated iron oxide nanoparticle (sIONP) loading following CPA loading, shown on an independent time axis. **c,** CPA unloading protocol, shown on an independent time axis. **d,** Representative cooling curve during vitrification; shading indicates the melting temperature (Tm) to glass transition temperature (Tg) window. Inset, cooling rates for biologically independent kidneys used to characterize the protocol (n = 13); each point represents one kidney and the black point and error bar indicate mean ± s.d. The dashed line indicates the critical cooling rate (CCR), below which ice formation is expected. **e,** Representative nanowarming curve. Inset, nanowarming rates for biologically independent kidneys (n = 10); we excluded three additional measurements obtained using a different method. Point and error-bar conventions are as in d. The dashed line indicates the critical warming rate (CWR), below which devitrification and ice formation are expected. **f, g,** Representative vitrified kidneys after CPA loading alone (f) or CPA and sIONP loading (g), shown by gross photography (top) and micro-computed tomography (microCT; bottom). The microCT scale is expressed in Hounsfield units (HU); red-orange (500–800 HU) indicates vitrified material, blue (<400 HU) indicates ice and dark red (>800 HU) indicates high sIONP concentration.

After PEZ42 loading, sIONPs (10 mg Fe/mL in PEZ42) were perfused via syringe pump at −15 °C (3–4 mL/min; mean pressure = 46.7 mmHg; n = 10) for 7–7.5 min (Fig. 2b). To prevent excessive iron accumulation in major vessels, which can induce localized hyperthermia during radiofrequency (RF) warming, the main renal artery and vein and the ureter were briefly flushed with a lower concentration sIONP solution (1.25 mg Fe/mL in PEZ42) before placing the organ in a cryobag prefilled with ∼100 mL of the 1.25 mg Fe/mL final solution.

### Vitrification, micro-CT characterization and nanowarming thermal performance

Thermal gradients between the surface and core risk thermomechanical cracking near the glass transition temperature (Tg ≈ −120 °C). To reduce thermal stress, we added an annealing step above the glass transition at −114 °C in a controlled-rate freezer (Fig. 2d). We optimized annealing duration to 100 min for ∼100-g kidneys, allowing temperature equalization with minimal ice nucleation or crystal growth. Organs were then cooled at 0.5 °C/min to −150 °C for cryogenic storage. Cooling rates in the kidney core, as measured by a temperature probe deep in the renal vein, averaged 3.15 ± 0.37 °C/min (n = 13), exceeding both the 2 °C/min design criterion and the CCR of PEZ42 (<1 °C/min; Methods) (Fig. 2d, inset).

During preliminary protocol development, we identified several failure modes associated with cracking and/or ice formation. These were mitigated through protocol refinements, including extending annealing duration; slowing the rate of cooling from the annealing temperature to below Tg; excluding kidneys with large, centrally located cysts or excessively high perfusion resistance; removing vascular and ureteral cannulas before vitrification; eliminating excess thermal mass, such as metal clamps used to secure temperature probes; and equilibrating sIONP Fe concentrations across the major vascular structures and the surrounding medium to approximate the average Fe concentration within the renal parenchyma. After these refinements, we observed no further cracking or ice formation.

We evaluated vitrification success visually and by cryogenic micro-CT (Fig. 2f,g) as we previously reported ^13^. Kidneys vitrified without sIONPs showed clear surrounding media and a stable kidney structure, consistent with a glassy state. sIONP-containing solutions are opaque, precluding visual assessment of ice (Fig. 2g). High-resolution scans confirmed uniform radiodensity (mean 735 HU) throughout cortical and medullary regions, with no visible cracking or ice crystallization.

Rewarming was performed in a 120-kW radiofrequency (RF) coil using a two-stage protocol (Fig. 2e). Stage 1 applied 50% RF power for 90 s, warming the organ from −150 °C to −120 °C at 22.8 ± 1.4 °C/min. Stage 2 increased power to 95%, rapidly heating the tissue through the devitrification zone to −10 °C at 32.3 ± 5.4 °C/min (total rewarming time: 340–350 s) (Fig. 2e, inset). This rewarming rate was 14-fold higher than the CWR of 92% PEZ42 (2.3 °C/min), the least stable concentration expected within the kidney. CPA was then unloaded on the circuit using a reverse multithermic protocol, in which temperature was raised from −8 °C to 4 °C as CPA concentration was stepped down, after which the kidney was flushed with LM6 carrier and then HTK cold storage solution (Fig. 2c).

### Perfusion and excretory recovery during normothermic machine perfusion

Following unloading, functional recovery was evaluated over 120 min of red blood cell-based NMP at 37 °C (Extended Data Figs. 1 and 2). VNW kidneys (n = 8) were compared with fresh controls (n = 6), 24-h SCS kidneys (n = 9) and CPA-only controls (n = 9). We stored VNW kidneys at −150 °C for 1–366 days; most were stored for 1–7 days.

Baseline donor characteristics (e.g., kidney weight) were comparable across groups, except for protocol-driven features such as cold ischemia time and for kidney laterality (Extended Data Table 1). Following NMP, CPA-only kidneys showed significantly greater weight gain from baseline (30.3 ± 11.1 g, 29.7 ± 11.0%) than VNW kidneys (11.0 ± 12.6 g, 11.0 ± 12.4%, P = 0.032 and 0.024, respectively), potentially indicating greater capillary leakage and edema in the CPA-only cohort. Weight gain in VNW kidneys was not statistically different from that of the fresh control or SCS groups.

During NMP, VNW kidneys showed prompt, uniform reperfusion and immediate urine production (Fig. 3a, Supplementary Video 1). Vascular and excretory function remained impaired relative to fresh and 24-h SCS kidneys but was similar to or numerically better than that of CPA-only controls (Fig. 3, Extended Data Table 2). Steady-state renal blood flow was 81.7 ± 21.5 mL/min/100 g after VNW, approximately 33% higher than after CPA exposure alone (61.6 ± 32.8 mL/min/100 g), although this difference was not statistically significant (P = 0.132; Fig. 3b,c). Correspondingly, intrarenal resistance was approximately 40% lower after VNW than after CPA exposure alone (1.04 ± 0.32 versus 1.73 ± 0.97 mmHg·min·100 g/mL; P = 0.043; Fig. 3d,e).

**Fig. 3.**
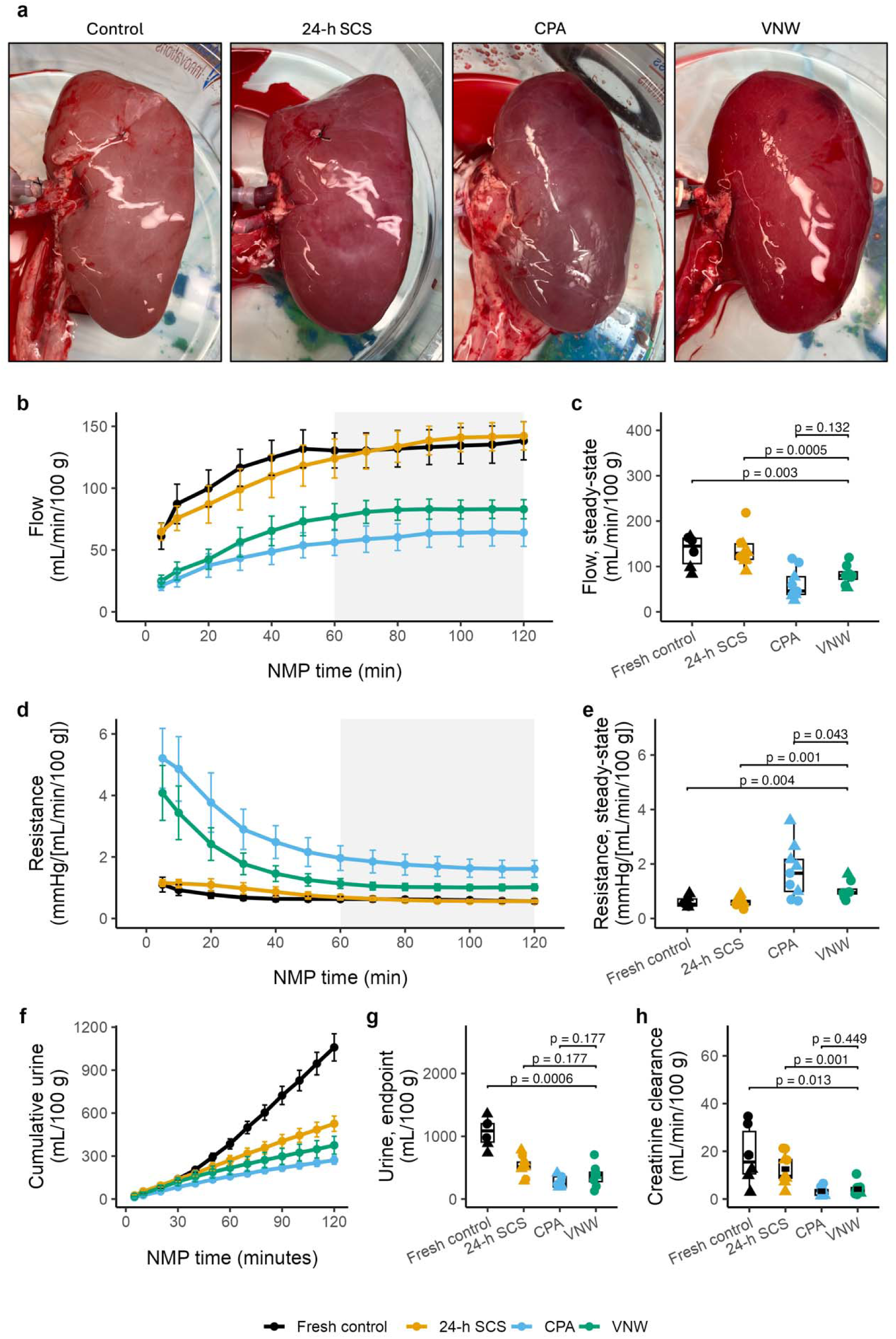
Vitrified and nanowarmed kidneys recover perfusion and excretory function. Vitrified and nanowarmed (VNW) kidneys consistently reperfused during normothermic machine perfusion (NMP) and produced urine, although renal blood flow and creatinine clearance remained reduced relative to fresh and 24-h static cold storage (SCS) kidneys. CPA-only kidneys underwent cryoprotective agent (CPA) loading and unloading without vitrification, cryogenic storage or nanowarming. **a,** Representative appearance of kidneys after reperfusion during NMP. **b,** Normalized renal blood flow over time. **c,** Steady-state renal blood flow, calculated as the mean from 60–120 min. **d,** Normalized intrarenal resistance over time. **e,** Steady-state intrarenal resistance, calculated as the mean from 60–120 min. **f,** Cumulative urine production over time. **g,** Cumulative urine production at 120 min. **h,** Creatinine clearance, calculated as the mean from 30–60 min. Biologically independent kidneys: fresh control, n = 6; 24-h SCS, n = 9; CPA-only, n = 9; VNW, n = 8. Time-course data are shown as mean ± s.e.m.; endpoint data are shown as box-and-jitter plots as described in Methods. Brackets indicate prespecified comparisons of VNW with each other group. P values were calculated using generalized least-squares models allowing group-specific variance, followed by estimated marginal means with two-sided tests and Holm adjustment for the three VNW comparisons; exact P values are shown.

VNW kidneys produced 375.6 ± 176.3 mL urine per 100 g over 120 min, approximately 38% more than CPA-only kidneys (271.3 ± 85.1 mL per 100 g) and 29% less than 24-h SCS kidneys (525.7 ± 160.2 mL per 100 g); neither comparison reached statistical significance after adjustment (both P = 0.177; Fig. 3f,g). Steady-state urine production showed a similar pattern (Extended Data Fig. 3b). Creatinine clearance was 4.16 ± 2.79 mL/min/100 g after VNW, 27% higher than after CPA exposure alone (3.28 ± 1.80 mL/min/100 g; P = 0.449), but remained below 24-h SCS (12.56 ± 6.43; P = 0.001) and fresh controls (18.35 ± 12.50; P = 0.013) (Fig. 3h, Supplementary Fig. 3).

### Metabolic profiles and biochemical injury markers

Metabolic activity was retained after vitrification and nanowarming, although recovery remained incomplete relative to fresh kidneys (Fig. 4, Extended Data Table 2). Steady-state oxygen consumption was 0.30 ± 0.08 mL/min/100 g in VNW kidneys, compared with 0.26 ± 0.14 mL/min/100 g in CPA-only kidneys and 0.37 ± 0.04 mL/min/100 g after 24-h SCS; VNW differed from fresh controls (0.47 ± 0.10 mL/min/100 g, P = 0.001) but not from CPA-only or SCS kidneys (Fig. 4a,b). Notably, venous lactate at 120 min was 0.90 ± 0.39 mmol/L after VNW, 41% lower than in CPA-only kidneys (1.53 ± 0.63 mmol/L) and 46% lower than after 24-h SCS (1.67 ± 0.81 mmol/L; P = 0.034 for both comparisons; Fig. 4c,d).

**Fig. 4.**
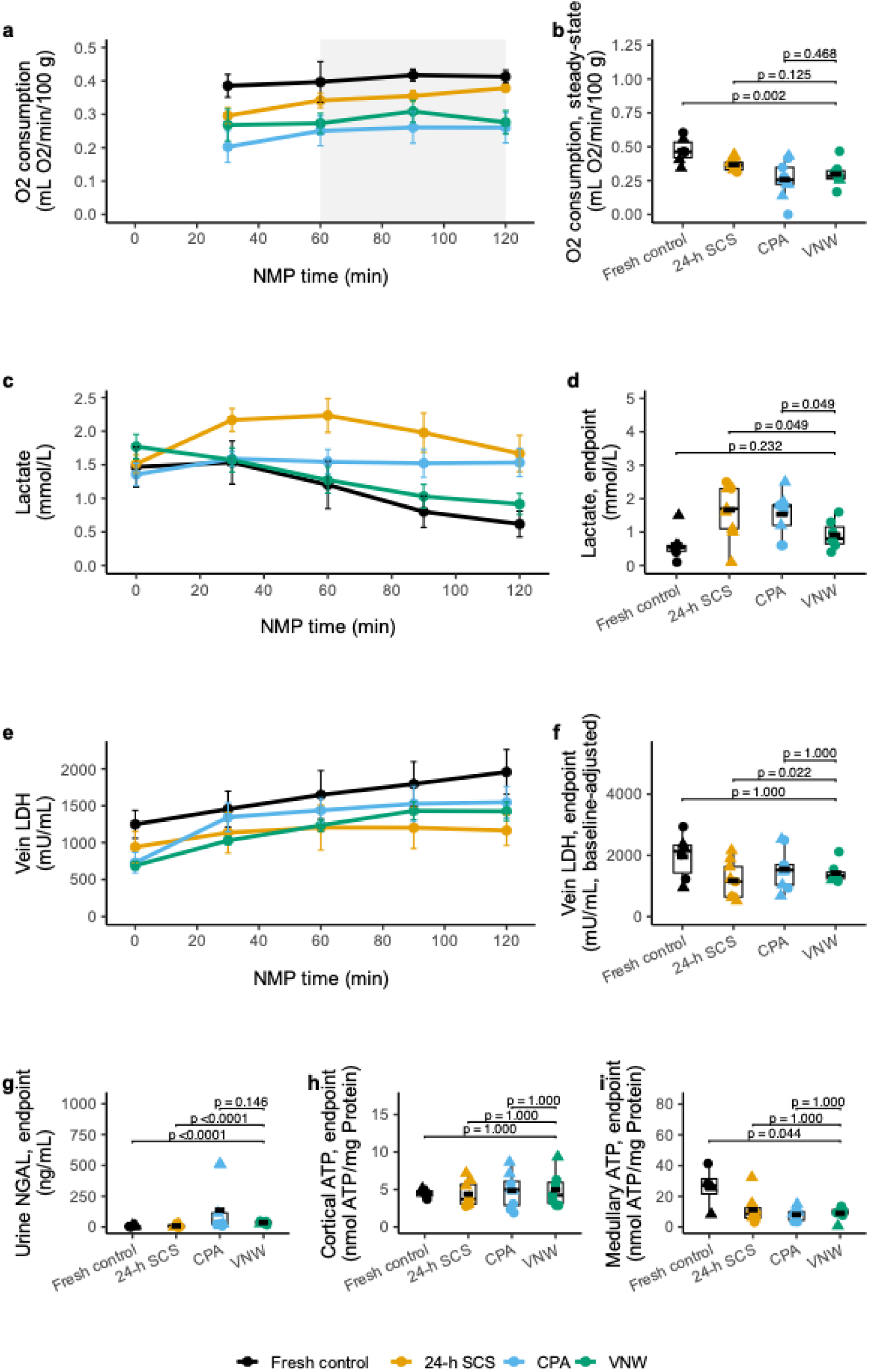
Metabolic and biochemical recovery after vitrification and nanowarming. Vitrified and nanowarmed (VNW) kidneys recover metabolic activity during normothermic machine perfusion (NMP), with residual differences from fresh and 24-h static cold storage (SCS) kidneys across some measures. Venous lactate was lower after VNW than in SCS and CPA-only kidneys. Cryoprotective agent (CPA)-only kidneys underwent CPA loading and unloading without vitrification, cryogenic storage or nanowarming. Box-and-jitter and trajectory plot conventions are as in Fig. 3. **a,** Oxygen consumption over time; n = 6, 9, 9 and 8 biologically independent kidneys for fresh, 24-h SCS, CPA-only and VNW groups, respectively. **b,** Steady-state oxygen consumption, calculated as the mean from 60–120 min; n as in a. **c,** Venous lactate over time; n as in a. **d,** Venous lactate at 120 min; n as in a. **e,** Venous lactate dehydrogenase (LDH) over time; n as in a. **f,** Venous LDH at 120 min, adjusted for the pre-NMP baseline; n as in a. Absolute change from baseline, a secondary measure, is shown in Supplementary Fig. 4. **g,** Urine neutrophil gelatinase-associated lipocalin (NGAL) at 120 min; n as in a. **h,** Cortical adenosine triphosphate (ATP) at 120 min; n = 4, 8, 8 and 7 biologically independent kidneys for fresh, 24-h SCS, CPA-only and VNW groups, respectively; fewer biopsies were available for this assay. **i,** Medullary ATP at 120 min; n as in h. For b, d, g, h and i, brackets show prespecified comparisons of VNW with each other group. P values were calculated using a generalized least-squares (GLS) model allowing group-specific variance, followed by estimated marginal means with two-sided tests and Holm adjustment for the three comparisons. For f, a GLS analysis of covariance (ANCOVA), including pre-NMP LDH as a covariate, was used with the same post hoc comparisons and adjustment. Exact P values are shown above the brackets.

Biochemical markers nevertheless demonstrated residual cellular and tubular injury. Baseline-adjusted terminal venous lactate dehydrogenase (LDH, a marker of cellular injury) was higher in VNW kidneys than after 24-h SCS but did not differ from fresh or CPA-only controls (Fig. 4e,f; Supplementary Fig. 4). Urinary neutrophil gelatinase-associated lipocalin (NGAL, a marker of tubular injury) at 120 min was elevated after VNW relative to fresh and 24-h SCS kidneys. However, NGAL was 73% lower after VNW than after CPA exposure alone (37.25 ± 10.84 versus 138.4 ± 212.2 ng/mL), although this difference was not statistically significant (P = 0.154; Fig. 4g).

Cortical adenosine triphosphate (ATP, a measure of cellular energetics) was preserved across groups, with mean values ranging from 4.40 to 4.96 nmol ATP/mg protein (Fig. 4h). In contrast, medullary ATP was lower after VNW than in fresh kidneys (8.94 ± 4.09 versus 25.91 ± 13.56 nmol ATP/mg protein; P = 0.044), but was similar to both 24-h SCS (11.35 ± 9.44 nmol ATP/mg protein) and CPA-only kidneys (8.08 ± 4.02 nmol ATP/mg protein) (Fig. 4i). Across 12 perfusate and urine chemistry parameters (six analytes, each in venous perfusate and urine), VNW kidneys did not differ significantly from CPA-only controls (all P ≥ 0.071; Supplementary Figs. 5 and 6).

### Histological injury and gross organ transplant suitability

Histological analysis showed that structural injury in VNW kidneys was modest (Fig. 5a). On the composite histologic injury scale (0–60; 12 features scored from 0 to 5), VNW kidneys had higher injury scores than fresh controls (11.62 ± 2.97 versus 6.50 ± 2.17; P = 0.012). VNW injury scores were not significantly different from those of 24-h SCS kidneys (8.22 ± 2.33; P = 0.091) or CPA-only controls (10.11 ± 2.76; P = 0.264; Fig. 5b). Subscore analysis showed that injury was concentrated predominantly within the tubular compartment, including acute tubular necrosis (ATN), tubular vacuolation and focal tubular epithelial injury (Fig. 5c; Supplementary Fig. 7). Glomerular and vascular injury remained minimal, with no evidence of widespread cortical necrosis. Trace amounts of dark staining were observed in scattered glomeruli and occasionally in tubules; Prussian blue staining was consistent with trace amounts of residual iron, as observed previously in rat kidneys ^13^.

**Fig. 5.**
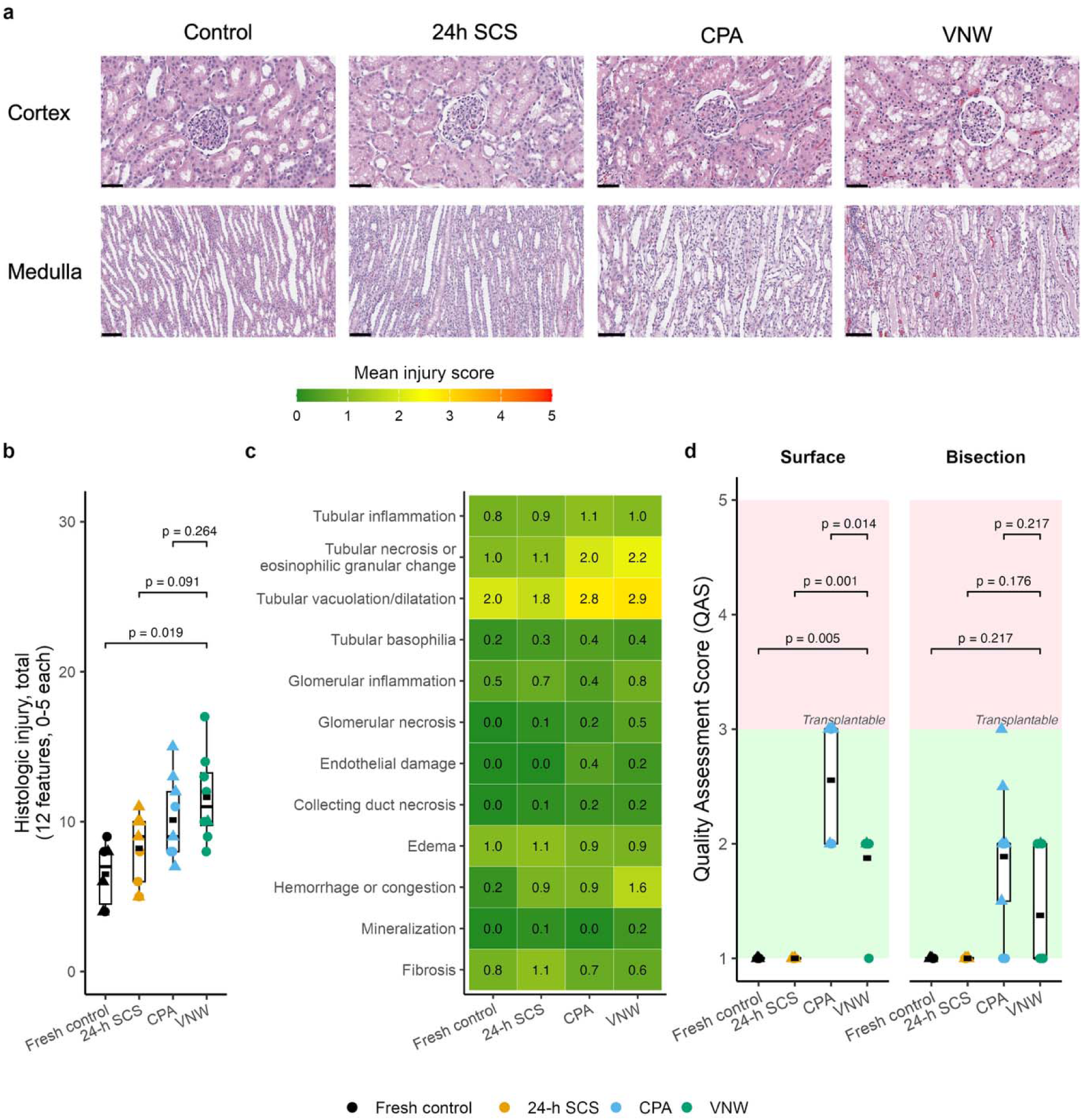
Vitrified and nanowarmed kidneys show predominantly tubular injury. Histological assessment of vitrified and nanowarmed (VNW) kidneys showed predominantly acute tubular injury, while gross transplant suitability remained preserved. Cryoprotective agent (CPA)-only kidneys underwent CPA loading and unloading without vitrification, cryogenic storage or nanowarming. Biologically independent kidneys per group: fresh control, n = 6; 24-h static cold storage (SCS), n = 9; CPA-only, n = 9; VNW, n = 8 (all panels). **a,** Representative cortical and medullary histology from each group. Scale bars, 50 µm (cortex) and 100 µm (medulla). **b,** Total histological injury score, calculated as the sum of 12 features scored from 0–5 (range, 0–60); box-and-jitter plot conventions are as in Fig. 3. Overall group effect: Kruskal–Wallis test, χ²(3) = 10.5, P = 0.015. Brackets show prespecified comparisons of VNW with each other group (two-sided Wilcoxon rank-sum tests, Holm-adjusted for the three comparisons and not conditional on the omnibus result); exact P values are shown above the brackets. **c,** Heat map of the 12 individual histological injury features by group; shading indicates the mean score for each feature (0–5). Data are descriptive and no statistical comparisons are shown. **d,** Hosgood Quality Assessment Scores based on surface (left) and bisection (right) assessments (1–5 scale); the dashed line indicates the transplant threshold (total score ≤3). Statistics as in **b**, applied separately to each assessment (Kruskal–Wallis: surface, χ²(3) = 26.9, P < 0.0001; bisection, χ²(3) = 15.4, P = 0.002). Box-and-jitter plot conventions are as in Fig. 3.

Gross viability was evaluated using the Quality Assessment Score (QAS) developed by Hosgood et al., in which scores of 1–3 are considered suitable for transplantation and scores ≥4 indicate poor organ quality ^18,19^. Standard surface assessment of perfusion quality yielded higher (worse) QAS values for VNW kidneys (1.88 ± 0.35) than for fresh (1.00 ± 0.00, P = 0.005) or SCS kidneys (1.00 ± 0.00, P = 0.001), but lower (better) values than for CPA-only controls (2.56 ± 0.53; P = 0.014; Fig. 5d, left). In some CPA-exposed kidneys, particularly on the dependent surface, superficial erythema adversely affected the macroscopic perfusion component of the QAS. After capsular removal and coronal bisection, this discoloration was confined to the superficial subcapsular tissue, whereas the underlying cortex appeared well perfused and similar to fresh controls (Extended Data Fig. 4, Supplementary Fig. 8). The localization of erythema to dependent surfaces suggested that it resulted from direct capsular contact with high-concentration CPA rather than impaired parenchymal perfusion. Accordingly, QAS values improved across CPA-exposed groups when macroscopic perfusion was scored from the internal cortical parenchyma appearance after kidney bisection rather than from the surface (Fig. 5d, right). All 32 study kidneys remained within the QAS range considered suitable for transplantation.

### Function after extended cryogenic storage

Although most VNW kidneys were stored for 1–7 days before nanowarming, one was stored for 366 days to test extended storage. After nanowarming, its function during NMP recovered within the range of the VNW group, with renal blood flow (80 mL/min/100 g) matching the VNW group mean, although urine production (129 mL/100 g/120 min) was lower than in most VNW kidneys. The lower urine production may have reflected higher baseline vascular resistance, as its initial vascular resistance during hypothermic perfusion before CPA loading was 2.31 mmHg·min·100 g/mL, compared with 1.73 ± 0.81 mmHg·min·100 g/mL for the rest of the group. Oxygen consumption (0.29 mL O /min/100 g) and venous lactate (0.80 mmol/L) were similar to the rest of the VNW group (0.30 ± 0.09 mL O /min/100 g and 0.91 ± 0.42 mmol/L, respectively). Vein LDH (1,656 mU/mL) and urine NGAL (50 ng/mL) were modestly higher than the rest of the group (1,425 ± 330 mU/mL and 35 ± 10 ng/mL), though LDH remained within the group’s overall range. Cortical and medullary architecture was well preserved on both gross and histologic examination (Supplementary Fig. 9), with a QAS equal to that of fresh controls (QAS = 1) and a composite histologic injury score (8 of 60) lower than that of the rest of the VNW group (12.14 ± 2.79).

### In vivo feasibility in an acute allotransplantation model

We evaluated surgical feasibility, reperfusion and immediate function in an acute, non-survival porcine allotransplantation model (Fig. 6). VNW, fresh and 24-h SCS kidneys were transplanted orthotopically (n = 3 per group). Following vascular anastomosis and clamp release, VNW grafts rapidly and uniformly reperfused, with restoration of normal gross color and appearance (Fig. 6a; Supplementary Video 2). Color Doppler ultrasound demonstrated arterial and venous perfusion throughout the renal parenchyma, and spectral Doppler showed preserved forward diastolic arterial flow and a low-resistance arterial waveform consistent with normal post-transplant graft perfusion (Fig. 6a).

**Fig. 6.**
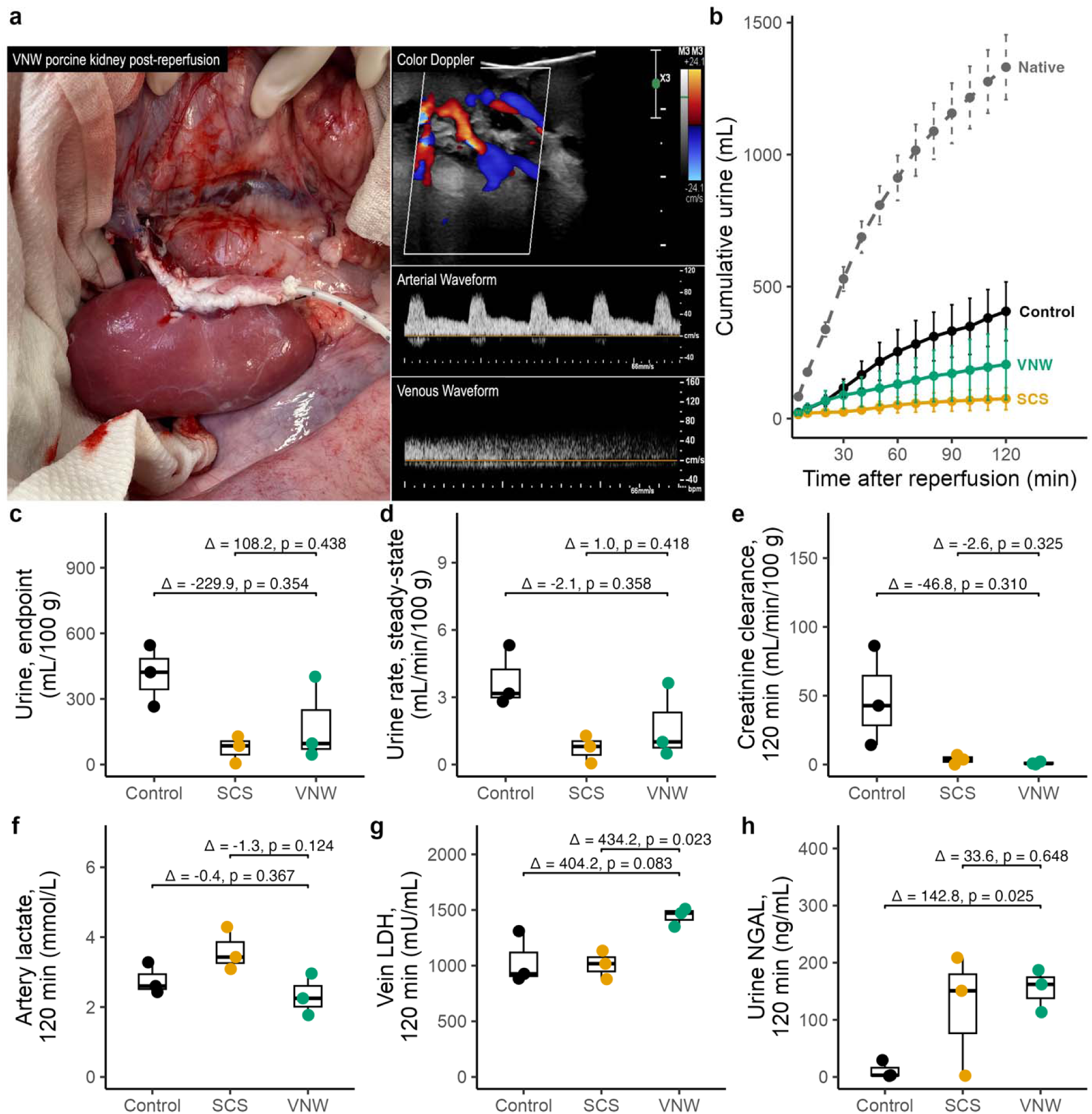
Nanowarmed kidneys rapidly reperfuse and sustain urine production after transplantation. Acute non-survival allotransplantation demonstrates the technical feasibility of transplantation and immediate in vivo function following vitrification and nanowarming (VNW). The transplant cohort comprised biologically independent kidneys from fresh control (n = 3), 24-h static cold storage (SCS; n = 3) and VNW (n = 3) groups. **a**, Gross appearance of a VNW kidney immediately following reperfusion, with color Doppler ultrasound demonstrating arterial and venous perfusion throughout the renal parenchyma and spectral Doppler waveforms demonstrating renal arterial and venous flow. **b**, Cumulative urine production over 120 min after reperfusion; native contralateral kidneys (n = 9) were pooled across all three groups and are shown as a reference. Points and lines indicate group means and error bars indicate ± s.e.m. **c**, Cumulative urine production at 120 min, normalized to kidney weight. **d**, Steady-state urine production rate, calculated as the mean from 60–120 min. **e**, Creatinine clearance at 120 min. **f**, Artery lactate at 120 min. **g**, Vein lactate dehydrogenase (LDH) at 120 min. **h**, Urine neutrophil gelatinase-associated lipocalin (NGAL) at 120 min. In **c–h**, each point represents one biologically independent kidney; boxes show the median (center line) and interquartile range, and whiskers extend to the most extreme value within 1.5 × the interquartile range. Δ denotes the raw mean difference between VNW and each comparator group (VNW minus comparator). P values were calculated using Welch’s two-sided t-test and Holm-adjusted for the two VNW comparisons within each panel; normality and equal-variance assumptions could not be verified at n = 3 per group.

VNW grafts produced urine immediately after reperfusion and sustained urine production throughout the 120-min observation period (Fig. 6b). Because the feasibility cohort was small, we could not verify the assumption of normality; in vivo comparisons are therefore descriptive and exploratory, and the study was not powered to detect between-group differences. Cumulative urine production was 181 ± 193 mL/100 g (range, 45–402) in VNW grafts, compared with 73 ± 62 mL/100 g (range, 5–128) after 24-h SCS and 411 ± 141 mL/100 g (range, 265–546) in fresh control grafts (Fig. 6c). Thus, mean urine production after VNW was approximately 2.5-fold greater than after 24-h SCS, although it remained lower than in fresh grafts and contralateral native kidneys (797 ± 165 mL/100 g; range, 587–1,039). The steady-state urine production rate (60–120 min) showed the same pattern: 1.7 ± 1.7 mL/min/100 g in VNW grafts, compared with 0.7 ± 0.6 after 24-h SCS and 3.8 ± 1.4 in fresh controls (Fig. 6d).

We next compared creatinine clearance and markers of metabolic stress and injury at 120 min (Fig. 6e–h). Overall, VNW grafts showed a pattern broadly similar to that of 24-h SCS grafts: numerically more urine output and lower lactate, similar creatinine clearance and urine NGAL, and higher LDH. Creatinine clearance was low in both preserved groups (VNW, 1.0 ± 1.0 mL/min/100 g; 24-h SCS, 3.6 ± 3.4) and well below that of fresh grafts (47.8 ± 36.3; Fig. 6e). Urine NGAL was similarly elevated in VNW (154 ± 37 ng/mL) and 24-h SCS (121 ± 107) grafts relative to fresh grafts (11 ± 16; Fig. 6h); the difference between VNW and fresh grafts was the only NGAL comparison that reached significance (Holm-adjusted P = 0.025). Arterial lactate was lower in VNW grafts (2.3 ± 0.6 mmol/L) than in 24-h SCS grafts (3.6 ± 0.6) and comparable to fresh controls (2.8 ± 0.4; Fig. 6f). In contrast, blood LDH was higher in VNW grafts (1,444 ± 84 mU/mL) than in 24-h SCS grafts (1,010 ± 128; Δ = 434 mU/mL; Holm-adjusted P = 0.023) and fresh controls (1,040 ± 235; Δ = 404 mU/mL; P = 0.083; Fig. 6g). No other VNW versus 24-h SCS comparison was significant (all Holm-adjusted P ≥ 0.12).

By the end of the 120-min observation period, VNW grafts had developed patchy ecchymosis that was predominantly subcapsular and resembled that observed after NMP (Extended Data Fig. 5; Supplementary Fig. 8). On bisection, some VNW grafts showed purplish discoloration of the medullary pyramids; similar, though somewhat less pronounced, medullary discoloration was also observed in some SCS and CPA-only kidneys after NMP. Histologic cortical and medullary injury, including endothelial injury and hemorrhage, appeared greater in transplanted VNW grafts than in transplanted fresh and SCS grafts or in VNW kidneys after NMP (Supplementary Figs. 10–11); these changes developed while urine production was sustained, although this small cohort was not powered for statistical comparison.

## Discussion

This study demonstrates vitrification, cryogenic storage for up to one year and nanowarming of ∼100-g porcine kidneys, followed by recovery of vascular, metabolic and excretory function ex vivo and immediate function after transplantation. Recovery remained incomplete relative to 24-h SCS, particularly for glomerular filtration and biochemical markers of tubular injury. However, despite exposure to cooling, vitrification, prolonged cryogenic storage and nanowarming, in addition to CPA loading and unloading, VNW kidneys showed no consistent functional decrement relative to kidneys exposed to CPA alone. Indeed, renal resistance and lactate were lower after VNW, while flow, urine production and creatinine clearance were numerically higher. These findings identify CPA-associated injury as a major remaining limitation while demonstrating that the physical processes required for long-term cryogenic storage can be scaled to human-sized kidneys while preserving substantial organ function.

Cryogenic organ banking could overcome the narrow time limits of hypothermic preservation by decoupling organ recovery from transplantation, thereby providing additional time for donor-recipient matching, organ assessment, recipient preparation, therapeutic intervention and broader geographic sharing ^2^. Efforts to achieve this goal have progressed from conventional freezing to vitrification, which avoids the structural injury associated with extensive ice formation ^3^. Early renal studies established tolerance to vitrifiable CPA concentrations and ultimately demonstrated the transplantation of a vitrified rabbit kidney, while identifying incomplete medullary CPA equilibration, toxicity, devitrification and thermomechanical stress as major remaining barriers ^5,20^.

Volumetric electromagnetic warming has addressed the principal physical barriers to organ rewarming by generating heat throughout the specimen rather than relying on conduction from the surface. Nanowarming uses magnetic nanoparticles distributed within the vasculature to generate heat in an alternating electromagnetic RF field ^10,11^. This approach was demonstrated in intact rat organs ^12,21–23^, enabling long-term cryogenic storage and life-sustaining transplantation of vitrified rat kidneys ^13^. An alternative approach, dielectric warming, directly couples radiofrequency electric fields to water and CPA molecules throughout the organ and has also achieved rapid volumetric warming of vitrified kidneys ^24–26^. Recent studies using 27- and 55-MHz dielectric warming demonstrated successful rewarming and transplantation of vitrified rabbit kidneys, including long-term survival with recovery of normal renal function after transplantation of a ∼14-g kidney ^25,26^. Together, these complementary approaches demonstrate that volumetric electromagnetic warming can overcome major limitations of surface warming and support functional recovery after organ vitrification. The present study extends this progress to ∼100-g porcine kidneys, approximately sevenfold larger than the ∼14-g rabbit kidneys and 100-fold larger than the rat kidneys previously recovered after vitrification, and to storage durations of up to one year, demonstrating functional recovery after an integrated VNW protocol at human organ scale.

Scaling organ vitrification to human dimensions shifts the challenge from rewarming alone to the integrated problems of cooling, CPA stability, CPA transport and biological toxicity. Heat-transfer limitations reduce achievable cooling rates as organ size increases ^14^, and the VMP formulation used for rat kidney vitrification does not provide sufficient stability at the slower cooling rates encountered in large kidneys, particularly in incompletely equilibrated tissue ^15^. Conversely, highly stable formulations such as M22 ^3,5^ require exposure to high CPA concentrations and showed excessive toxicity under the conditions evaluated in our preliminary studies. Thus, although volumetric warming can overcome a major historical barrier to organ vitrification, successful preservation at the human-organ scale also requires CPA formulations and delivery protocols compatible with the slower cooling rates and longer transport distances imposed by larger organs.

We therefore developed an integrated protocol tailored to these thermal and transport constraints. PEZ42 was formulated to provide vitrification stability at the cooling and warming rates achievable in porcine kidneys while minimizing CPA exposure. Multithermic perfusion further limited high-concentration CPA exposure at warmer temperatures, reflecting the strong temperature dependence of CPA toxicity and the need to balance tissue equilibration with cumulative exposure ^3,16,27^. Controlled cooling with an annealing step at −114 °C reduced thermomechanical stress and, together with refinements to organ selection and handling, prevented detectable cracking and ice formation. Two-stage RF nanowarming subsequently achieved warming rates well above the measured critical warming rate of PEZ42.

CPA distribution within the renal medulla remains particularly important because renal vascular anatomy limits delivery to this region. Historical studies of rabbit kidneys identified incomplete medullary equilibration as a major determinant of vitrification failure ^5^. In our prior study, modeling predicted that medullary tissue reached approximately 92% of the nominal perfusate concentration ^15,16^, while DSC measurements indicated that PEZ42 at this concentration remained compatible with the cooling and warming rates achieved experimentally. These results underscore the importance of considering CPA transport and thermal requirements together rather than treating formulation, loading and heat transfer as independent problems. Sustained vascular perfusion, oxygen consumption and urine production during NMP provide complementary biological evidence against catastrophic medullary injury, although lower medullary ATP than in fresh kidneys indicates that medullary injury was not entirely avoided.

Comparison with CPA-only controls provides important insight into the origin of residual biological injury. Across renal blood flow, intrarenal resistance, urine production, creatinine clearance, oxygen consumption and multiple metabolic and biochemical measures, VNW kidneys showed no detectable functional decrement relative to CPA-only kidneys. This is notable because VNW kidneys underwent the same CPA loading and unloading, followed by the additional stresses of vitrification, cryogenic storage and nanowarming. VNW kidneys also had lower venous lactate than CPA-only kidneys and lower (better) QAS values. Taken together, these findings suggest that CPA exposure and delivery are the predominant sources of residual functional injury and that the subsequent cryogenic steps add little detectable functional or metabolic injury under the conditions tested. This interpretation is consistent with earlier small-animal studies in which nanowarmed organs showed biological outcomes similar to CPA-treated controls ^12,21,22^. Histological injury was numerically greater in VNW than CPA-only kidneys, although this difference was not statistically significant, indicating that additional structural injury from the cryogenic process cannot be excluded. Nevertheless, the absence of a corresponding functional decrement suggests that the major opportunity to improve organ recovery now lies in further reduction of injury associated with CPA exposure and delivery.

Importantly, CPA toxicity is increasingly amenable to systematic optimization. Vitrification-solution toxicity depends strongly on CPA composition rather than total concentration alone ^3^. Mathematical modeling and high-throughput screening can identify CPA combinations that balance vitrification stability with specific, nonspecific and interaction-dependent toxicity ^27,28^. Quantitative transport modeling can optimize organ-specific loading and unloading ^16^. Biological strategies that modify cellular responses to concentrated CPAs offer an additional opportunity to reduce injury ^29^. Together, these approaches provide a rational path toward closing the remaining performance gap between VNW and conventional hypothermic preservation.

Histopathological injury after VNW was predominantly confined to the tubular compartment, whereas glomerular and vascular injury was comparatively limited, and composite injury scores did not differ significantly from those after 24-h SCS. This pattern may be favorable for recovery because surviving tubular epithelial cells retain substantial regenerative capacity after acute injury ^30^. However, reversibility cannot be determined from short-duration NMP, and structural injury may coexist with preserved global perfusion, urine production and oxygen consumption ^31^. Survival transplantation will be required to determine the long-term significance of the tubular injury observed here.

Small amounts of residual iron were identified focally in some VNW kidneys, consistent with our prior observations in rats ^13^. The sIONPs used for nanowarming were developed for high heating efficiency, colloidal stability, biocompatibility and vascular washout ^32^. Recent in vivo studies found no overt toxicity at doses comparable to or greater than those in a theoretical fully loaded organ, and estimated systemic exposure from residual iron after organ washout was orders of magnitude lower ^33^. Nevertheless, residual nanoparticle distribution, washout and long-term biological effects will require continued evaluation for clinical translation.

Acute allotransplantation demonstrated that vitrified and nanowarmed porcine kidneys reperfused uniformly, maintained vascular patency and produced urine immediately and throughout the 2-h observation period. Histologically, however, transplanted VNW grafts showed greater glomerular and endothelial injury, with parenchymal hemorrhage, than VNW kidneys assessed by NMP. Compared with transplanted SCS grafts, VNW kidneys showed somewhat greater urine production and lower arterial lactate, with similar creatinine clearance and urine NGAL, but higher LDH consistent with the greater histologic injury. Several conditions present in vivo but absent or attenuated during NMP may have contributed, including additional warm ischemia during vascular anastomosis, the systemic inflammatory response to major surgery, and exposure to circulating leukocytes, platelets, complement and other undiluted plasma factors. In contrast, NMP used diluted, leukocyte-depleted autologous blood supplemented with dexamethasone. Because the histologic injury was more pronounced in VNW grafts than in transplanted fresh and SCS grafts, the renal endothelium of VNW kidneys may be more susceptible to these additional insults. Given the similar ex vivo function of VNW and CPA-only kidneys, CPA exposure is a likely contributor, although CPA-only kidneys were not transplanted and the relative contributions of CPA exposure and the cryogenic steps remain to be defined. Identifying and mitigating this susceptibility will be an important focus of future work.

In our previous rat studies, VNW kidney dysfunction peaked 3–4 days after transplantation before recovering over approximately 2 weeks ^13^, making longer-term survival transplantation an important next step. We initially considered a 30-day porcine autotransplant model, but this would require three survival operations (kidney procurement, delayed autotransplantation and subsequent contralateral nephrectomy), with substantial surgical and logistical complexity. Conversely, survival allotransplantation would require immunosuppression and introduce alloimmune injury as an additional confounder. We therefore used acute allotransplantation to first establish that VNW kidneys could be surgically transplanted and demonstrate immediate vascular and excretory function in vivo. Longitudinal survival transplantation will now be required to define the trajectory of renal injury and recovery and determine whether VNW kidneys can provide sustained, life-supporting renal function.

Although most VNW kidneys in this study were stored for 1–7 days, one kidney was maintained in the vitrified state for 366 days before nanowarming and recovered function within the range of the VNW group, with renal blood flow similar to the group mean and a transplant suitability score (QAS) equal to that of fresh controls. This observation is consistent with our previous demonstration that vitrified rat kidneys could be stored cryogenically for up to 100 days before successful transplantation ^13^. We have similarly shown that vitrified pancreatic islets remain stable during extended cryogenic storage, with preserved function after transplantation following storage for at least 9 months ^34^ and up to 2 years in subsequent unpublished studies. Although the present study was not designed to determine the effect of storage duration on kidney recovery, these findings across organs and cellular tissues support the premise that, once successfully vitrified and maintained under appropriate cryogenic conditions, biological materials can remain stable for extended periods. Thus, the principal constraints on VNW organ preservation may ultimately be successful vitrification and rewarming rather than the duration of storage itself.

Several limitations warrant consideration. Group sizes were small, particularly for transplantation (n = 3 per group), and extended storage was tested in a single kidney; the absence of statistically significant differences should therefore not be interpreted as equivalence. Although porcine kidneys approximate human kidney size, differences in vascular anatomy, CPA transport and biological susceptibility will require validation in human kidneys declined for transplantation. Renal recovery was evaluated primarily during short-duration NMP and acute transplantation, and no validated set of NMP viability markers definitively predicts long-term post-transplant function ^31^. Survival transplantation will therefore be required to determine sustained glomerular filtration, recovery from tubular injury and chronic histological outcomes. Thermometry and imaging sample only portions of a spatially heterogeneous organ and cannot exclude microscopic ice formation at every location. Furthermore, although the similarity between VNW and CPA-only kidneys strongly implicates CPA exposure as the predominant source of residual dysfunction, the present design cannot exclude a smaller contribution from the cryogenic process, and CPA-only kidneys were not transplanted to determine whether CPA exposure alone accounts for the endothelial injury observed in vivo. Finally, VNW requires more complex perfusion, nanoparticle, cryogenic and RF systems than conventional preservation and will require further standardization, automation and validation before clinical implementation.

In conclusion, vitrification and volumetric nanowarming can be extended from rodent kidneys to human-scale porcine kidneys, with functional recovery after up to one year of cryogenic storage and immediate function after transplantation. The physical requirements for cooling and rewarming organs at this scale were met while preserving substantial vascular, metabolic and excretory function, with several outcomes approaching those observed after 24-h SCS. Importantly, VNW produced little detectable functional decrement beyond that attributable to CPA exposure alone, suggesting that CPA toxicity and delivery, rather than the physical processes of vitrification and nanowarming, are now the principal barriers to further improvement. Optimization of CPA composition and delivery, followed by survival transplantation and validation in human kidneys, provides a path toward clinically relevant organ cryobanking. Together, these findings establish long-term banking of human-scale organs as a tractable engineering and biological challenge and define the remaining barriers to its clinical translation.

## Methods

### CPA formulations and nanoparticle preparation

Two cryoprotective agent (CPA) solutions were used for organ perfusion: PMP and PEZ42. PMP contained 22.3% w/v dimethyl sulfoxide (DMSO; >99%, Sigma-Aldrich, USA), 16.8% w/v ethylene glycol (Fisher Scientific, USA), 10.9% w/v propylene glycol (>99%, Sigma-Aldrich, USA), 1% w/v X-1000 and 1% w/v Z-1000 in LM5 carrier solution (total concentration, 52.0% w/v; 7.01 M). PEZ42 contained 22.3% w/v DMSO, 20.8% w/v ethylene glycol, 12.9% w/v propylene glycol, 1% w/v X-1000 and 3% w/v Z-1000 in LM5 (total concentration, 60.0% w/v; 7.95 M). Synthetic ice blockers X-1000 and Z-1000 were obtained from 21st Century Medicine (Fontana, CA, USA).

Dynamic concentration ramps during PMP loading were generated by mixing full-strength PMP with LM6, a carrier solution consisting of LM5 ^3^ supplemented with 25 mM D-mannitol (99.9%, Chem-Impex, USA) to maintain osmolality. The hyperosmolar unloading solution (52% PMP + 275 mM mannitol) was prepared by adding D-mannitol to diluted PMP. Silica-coated iron oxide nanoparticles (sIONPs) were synthesized from EMG308 iron oxide core particles (Ferrotec, Santa Clara, CA, USA), coated with a protective silica shell and surface-modified with polyethylene glycol and chlorotrimethylsilane (TMS; >99%, Sigma-Aldrich, USA). The synthesized sIONPs were suspended in PEZ42 and filtered through a 0.22 µm membrane prior to use ^32,35^.

### Critical cooling and warming rates

Critical cooling and warming rates (CCR and CWR) were determined by differential scanning calorimetry (DSC), as previously described ^15^. For CWR determination, samples were vitrified at the maximum programmable cooling rate and then rewarmed at 1–6 °C/min (n = 3 per rate). The devitrification temperature (Td) was identified from the crystallization exotherm, and the melting temperature (Tm) was defined as the peak of the melting endotherm measured at 1 °C/min. Because Td increases approximately linearly with the logarithm of the warming rate, Td was plotted against log(warming rate) and fitted by linear regression. CWR was defined as the warming rate at which 0.2% ice crystallization occurs. CCR was assessed by cooling samples at 1 °C/min (n = 3). No detectable crystallization exotherm was observed during cooling in any replicate, establishing a CCR of <1 °C/min.

### Kidney procurement and initial flush

Castrated male Yorkshire crossbreed pigs (19–36 kg) served as kidney donors. All procedures were approved by the Institutional Animal Care and Use Committee (IACUC) at the University of Minnesota. Following midline laparotomy, the kidneys were mobilized, and the renal artery, renal vein and ureter were dissected. Intraoperative ultrasound was performed to screen for large (> 1.5 cm) or centrally located cysts. Systemic heparinization was achieved via intravenous administration (300 IU/kg). Five minutes post-administration, the renal artery was cannulated with a 2 mm vessel cannula (Medtronic DLP, REF 30004). The kidney was immediately flushed with room-temperature Lactated Ringer’s solution supplemented with heparin (5000 IU/L) and papaverine (60 mg/L) to prevent vascular spasm and thrombosis. After 2 min, the flush was switched to cold (4 °C) heparinized (1,000 IU/L) Histidine-Tryptophan-Ketoglutarate (HTK) solution (∼200 mL). The kidney was weighed and stored in HTK on ice during transit. For the initial development of the protocol, organ recovery was performed using a donation after circulatory death (DCD) model, in which heparinized donors were euthanized and, after confirmation of death, rapid organ recovery was performed with minimal warm ischemia time. We then switched to a donation after brain death (DBD) model, in which organs were recovered during a terminal anesthetic procedure. Because retrospective comparison showed no differences between DCD and DBD kidneys, data were pooled for reporting. Throughout the figures, DBD kidneys are presented as circles and DCD as triangles.

### Multithermic CPA loading protocol

Kidneys were connected to a custom-built, pressure- and flow-regulated open-loop multithermic machine perfusion system ^13^. Initial perfusion was conducted at 40 mmHg for 5 min using LM6 to stabilize flow and establish baseline vascular resistance. CPA loading was initiated by ramping PMP from 0 to 4.14 M at 51.75 mM/min over 80 min at 40 mmHg. Following a 5 min equilibration hold at 4.14 M, perfusion pressure was ramped from 40 to 60 mmHg while PMP was increased to 7.01 M (full-strength) over 5 min.

Upon reaching full-strength PMP, perfusate and ambient chamber temperatures were rapidly lowered to below −15 °C. Perfusion was maintained at −15 °C and 60 mmHg for 10–15 min. The perfusate was then transitioned to PEZ42 (7.95 M), and perfusion pressure was increased to 80 mmHg over 5 min to compensate for temperature-induced viscosity changes. Full-strength PEZ42 perfusion was maintained at −15 °C for 15–20 min to ensure tissue equilibration. Peak CPA perfusion was monitored dynamically by measuring the refractive index of ureteral effluents and stopping CPA loading when the effluent reached 97.5% of full strength.

### Nanoparticle perfusion

Following PEZ42 loading, kidneys were transitioned to a syringe-pump circuit maintained at −15 °C for sIONP loading. A suspension of sIONPs (10 mg Fe/mL) in PEZ42 was perfused at a constant flow rate of 3–4 mL/min for 7–7.5 min while monitoring line pressure. To reduce high nanoparticle concentrations in the major vessels while maintaining sIONP distribution in the microvasculature, the main arterial and venous trunks were subsequently flushed with 3–4 mL of PEZ42 containing 1.25 mg Fe/mL sIONPs, and non-flushed channels were selectively occluded. The ureter and renal pelvis were similarly flushed with 5 mL of the 1.25 mg Fe/mL sIONP solution to displace residual fluid and eliminate voids.

The kidney was placed in a 2-mil polyethylene bag (Uline, Pleasant Prairie, WI, USA) prefilled with 100–110 mL of matching solution (1.25 mg Fe/mL sIONPs in PEZ42). For thermal characterization cohorts, fiberoptic temperature probes (Micronor Sensors, Ventura, CA, USA) were positioned in the surrounding fluid and inserted directly into the renal core via the renal vein. For functional evaluation and transplantation cohorts, a single fiberoptic probe was placed in the adjacent fluid to monitor thermal history at 1 s intervals without compromising parenchymal integrity.

### Vitrification and cryogenic storage

Vitrification was performed in a controlled-rate freezer (Kryo 560, Planer Ltd., Middlesex, UK). The chamber was pre-cooled to −20 °C and ramped to −114 °C at −40 °C/min. An isothermal annealing step was held at −114 °C for 100 min (slightly above the glass transition temperature, Tg ≈ −120 °C) to promote thermal equilibration across the organ mass while mitigating thermomechanical stress, as cracking during cooling and/or cryogenic storage was the primary physical failure mode observed during protocol development, particularly at larger vitrified volumes. Following annealing, the chamber was cooled at −0.5 °C/min to −150 °C to safely transition the organ into the glassy state. After an isothermal hold at −150 °C for 20–40 min, the vitrified kidney was transferred to a cryogenic storage freezer (PHC Corporation of North America, Wood Dale, IL, USA) and maintained at −150 °C for 1 day to 1 year.

### Nanowarming and post-rewarming CPA unloading

Nanowarming was conducted using a 120-kW 360 kHz radiofrequency (RF) induction coil (AMF Life Systems, Auburn Hills, MI, USA) operating under a two-stage rewarming protocol ^14^. The kidney was placed in the RF coil and exposed to 50% power (18.36 kA/m) for 90 s, warming the organ to approximately −120 °C. Coil output was then increased to 95% power (33.33 kA/m) to rapidly drive the sample through the critical warming phase to between −20 °C and −10 °C (total exposure: ∼350 s).

Following nanowarming, the organ was immediately transferred to the perfusion system to initiate CPA removal within 2 min. Unloading was initiated with full-strength PMP at −8 °C for 5 min, followed by 52% PMP (3.65 M) supplemented with 275 mM mannitol at −8 °C for 15 min at 40 mmHg. Over the subsequent 120 min, the perfusate concentration was linearly reduced from 3.65 M to 0 M, while the temperature was stepped to −3 °C for 10 min and then maintained at 0–4 °C. Finally, organs were flushed with LM6 for 15–30 min, followed by HTK for 10–15 min at 40 mmHg. Unloaded kidneys were stored in ice-cold HTK prior to evaluation.

### Micro-CT imaging and radiodensity analysis

Micro-computed tomography (micro-CT) was performed at a spatial resolution of 0.098 mm using an XT H 225 system (Nikon Metrology, MI, USA) ^13^. Vitrified kidneys enclosed in cryobags were held under liquid nitrogen vapor (−150 °C) in an insulated container during scanning (121 kV, 150 µA). Water and air reference tubes at ambient temperature were scanned concurrently for Hounsfield unit (HU) calibration. Reconstructions were performed using 3D CT Pro (Nikon Metrology) with beam-hardening corrections, converted to 16-bit format and then processed in VGStudio Max 3.2 (Volume Graphics, NC, USA) and MATLAB (MathWorks, Natick, MA, USA) to calculate spatial radiodensity across cortical and medullary regions.

### Normothermic machine perfusion (NMP) functional assessment

Organ function was evaluated using an ex vivo NMP circuit comprising a perfusate reservoir, a peristaltic roller pump (MasterFlex, Cole-Parmer, Vernon Hills, IL, USA), a bubble trap, a membrane oxygenator (Medtronic, Minneapolis, MN, USA), an in-line arterial filter (Medtronic, Minneapolis, MN, USA), pressure sensors (PendoTech, Princeton, NJ, USA) and heat exchangers maintained at 37 °C (Extended Data Fig. 2). The perfusate consisted of 300 mL of autologous, leukocyte-depleted whole blood, 130 mL of Williams’ Medium E without phenol red, 47.5 mg creatinine, 12 mg mannitol, 4.8 mg dexamethasone, 6 mg mixed amino acids, 0.25 mL multivitamins, 5 IU regular insulin, 1000 IU heparin and 5 mg verapamil. Continuous infusions of multivitamins (0.3 mL/h), amino acids (30 mg/h), insulin (18 IU/h) and verapamil (1.5 mg/h) were delivered via syringe pump (New Era Pump Systems, Farmingdale, NY, USA). Electrolyte levels and pH were maintained with supplemental 8.4% sodium bicarbonate, 5% dextrose, potassium chloride (2 mEq/mL) and calcium gluconate (100 mg/mL).

Perfusion was maintained at a target mean pressure of 75 mmHg (up to 100 mmHg for treated groups) for 120 min. Intrarenal vascular resistance (IRR) was computed as:

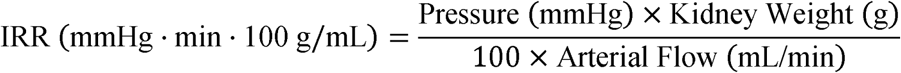

Urine output was continuously measured, and volume was replaced 1:1 with a 2:1ratio of Plasma-Lyte (Baxter Healthcare Corporation, IL, USA) and sterile water (ICU Medical Inc, IL, USA). Blood gases and electrolytes were analyzed every 30 min (ABL90 FLEX, Radiometer, Brea, CA, USA). Oxygen consumption rate (OCR) was calculated using the Fick principle, based on arterial-venous oxygen content differences and renal blood flow:

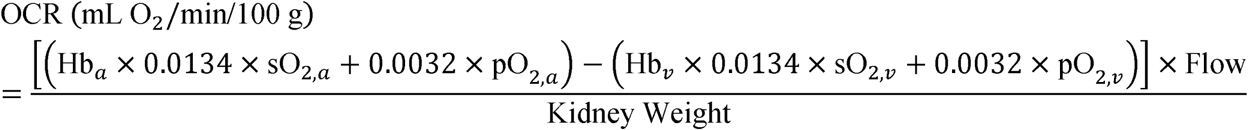

The timeclock for urine and perfusate sampling began 1 min after reperfusion to allow the kidney to warm to near 37 °C.

### Biochemical and biomarker assays

Venous perfusate and urine samples collected every 30 min were centrifuged (3,500, 10 min, 4 °C) and then stored at −80 °C. Creatinine in perfusate and urine was quantified using an enzymatic colorimetric assay (Crystal Chem, Elk Grove Village, IL, USA) on a Synergy 2 SL microplate reader (BioTek, Winooski, VT, USA). Creatinine clearance (CrCl) was calculated as:

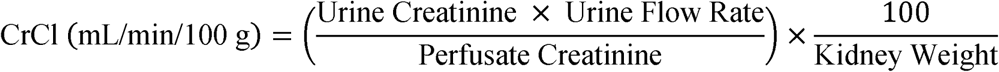

Tissue adenosine triphosphate (ATP) was quantified from 3 mm cortical and medullary biopsies (Miltex, Electron Microscopy Sciences, Hatfield, PA, USA) obtained immediately after NMP, flash-frozen in liquid nitrogen and homogenized in 70% ethanol containing 2 mM EDTA (pH 10.9). ATP was measured by luminescence (ATP Bioluminescence Assay Kit CLS II, Roche, Switzerland) and normalized to total protein content determined by the Bradford assay (Bio-Rad, Hercules, CA, USA).

Lactate dehydrogenase (LDH) activity in venous perfusate was quantified using a luminescent cytotoxicity assay (LDH-Glo, Promega, Madison, WI, USA). Urinary neutrophil gelatinase-associated lipocalin (NGAL) was measured by enzyme-linked immunosorbent assay (Pig NGAL ELISA Kit, Abcam, Cambridge, UK) using a 4-parameter logistic (4PL) regression curve.

### Histopathology and gross organ suitability scoring

Post-NMP kidneys were bisected along the coronal plane. Medullary and cortical wedge biopsies were fixed in 10% neutral buffered formalin, paraffin-embedded, sectioned and stained with hematoxylin and eosin (H&E). Whole-slide images were acquired with an Aperio GT450 scanner (Leica Biosystems, Vista, CA, USA) and evaluated by a senior, board-certified pathologist blinded to treatment arms (HistoWiz, Brooklyn, NY). Histologic damage was quantified using a semi-quantitative scoring system across 12 parameters (0–5 scale per feature): tubular inflammation, tubular necrosis/granular change, tubular vacuolation/dilation, tubular basophilia, glomerular inflammation/hematin deposition, glomerular necrosis, endothelial damage, collecting duct necrosis, edema, hemorrhage/congestion, mineralization and fibrosis (Supplementary Table 2).

Gross organ suitability was assessed using the Hosgood Quality Assessment Score (QAS) criteria, an ordinal scale on which each kidney receives a single integer score from 1 to 5 based on renal blood flow, urine production and macrovascular surface color (Supplementary Table 1); scores of 1–3 indicate transplantable quality and scores ≥4 indicate non-viability ^18^. To account for superficial CPA contact effects, gross perfusion scoring was also performed after parenchymal bisection to evaluate the internal cortical appearance (Fig. 5d).

### Non-survival kidney allotransplantation

Surgical feasibility and immediate post-reperfusion function were evaluated in a non-survival porcine allotransplantation model. Recipient pigs (60–80 kg) were anesthetized, intubated and prepared via midline laparotomy. Norepinephrine was infused through a central venous catheter to maintain systolic blood pressure above 70 mmHg throughout the procedure. The right native kidney was mobilized on its vascular pedicle, and the left native ureter was cannulated to monitor baseline function.

Systemic heparinization (100–200 IU/kg) and immunosuppressive/anti-inflammatory coverage with methylprednisolone (250 mg) were administered intravenously. The recipient right kidney vessels were clamped, and nephrectomy was performed. The donor kidney (fresh control, 24-h SCS or VNW) was revascularized by performing end-to-end arterial and venous anastomoses to the right renal vascular stumps with 6-0 polypropylene sutures for the vein and 7-0 for the artery. Prior to reperfusion, intravenous mannitol (12.5 g) and furosemide (0.5–0.8 mg/kg) were administered to promote diuresis. Vascular clamps were removed to initiate reperfusion.

Macrovascular surface color restoration, renal blood flow, systemic hemodynamic stability and urine production were monitored continuously throughout a 120-min post-reperfusion observation period. Arterial blood was sampled periodically, and urine was collected separately from both the donor organ and the contralateral native ureter. Additional heparin was administered post-reperfusion and re-dosed based on activated clotting time (ACT) measurements (total heparin dose 150–800 IU/kg). At the 120-min time point, the transplanted donor kidney and remaining native kidney were explanted, bisected and sampled (cortical and medullary tissue) for ATP quantification and histological analysis.

### Statistical analysis

Analyses were conducted in R 4.5.0 using standard statistical packages (nlme 3.1-168, lme4 1.1-37, lmerTest 3.1-3, emmeans 2.0-4, car 3.1-3, outliers 0.15). The kidney was the unit of analysis; repeated within-kidney measurements were not treated as independent. The main NMP cohort comprised 32 kidneys after the two exclusions described below (fresh control n = 6, 24-h SCS n = 9, CPA-only n = 9, VNW n = 8). One kidney was excluded because of extensive renal cysts. A 24-h SCS kidney was excluded from all analyses as a statistical outlier for steady-state arterial flow (426 vs. next highest 218 mL/min/100 g; two-sided Grubbs’ test, n = 10, G = 2.67, P = 0.0006); the significance of the VNW comparisons for steady-state flow and 120-min urine output was unchanged when this kidney was restored. Significance was set at P < 0.05, two-sided. Data are presented as mean ± SD unless otherwise stated. Where samples were unavailable for an endpoint, the number of kidneys analyzed is given in the corresponding figure legend.

Baseline and procurement characteristics (procurement type, kidney laterality, warm and cold ischemia times, kidney weight at each stage, and weight gain from procurement to the end of NMP, in grams and as a percentage) were compared using the three planned VNW-versus-comparator contrasts described below with Holm adjustment: Wilcoxon rank-sum for continuous variables and Fisher’s exact test for categorical variables. Warm ischemia time was determined by procurement type (5 min DBD, 10 min DCD), and cold ischemia time was fixed by experimental group (3–3.5 h fresh control, 24 h for 24-h SCS, 7 h for CPA-only and VNW), rather than varying independently within each arm. Therefore, between-group differences in these two variables reflect study design rather than uncontrolled procurement variability.

For continuous endpoints in the NMP cohort (flow, resistance, urine output and rate, creatinine clearance, oxygen consumption, perfusate and urine chemistry, venous lactate and bicarbonate, NGAL, ATP), between-group comparisons were limited to three planned contrasts, VNW against each other group, rather than all pairwise combinations. Steady-state values were the mean of measurements from 60–120 min (flow, resistance, urine production rate, oxygen consumption). Creatinine clearance was summarized as the mean of the 30- and 60-min values because perfusate creatinine declined progressively during perfusion (Supplementary Fig. 3). Cumulative urine, perfusate and urine chemistry (pH, sodium, potassium, chloride, calcium and glucose, each measured in venous perfusate and urine; 12 parameters in total), venous bicarbonate and lactate, urine NGAL and ATP were analyzed at 120 min. Each endpoint was fit by generalized least squares with group as the fixed effect and a group-specific residual variance structure. Log scaling was screened for flow, resistance and urine output, and the raw scale was retained for all reported endpoints. Urine NGAL is right-skewed in the CPA-only group (two of nine kidneys near 510 ng/mL; median 23.4 ng/mL), so we retained the raw scale with group-specific variances and confirmed the VNW contrasts with Holm-adjusted Wilcoxon rank-sum tests and a log(x+1) model; conclusions were unchanged (VNW vs fresh control and 24-h SCS, P ≤ 0.005 by rank test; VNW vs CPA-only, P = 0.47). The omnibus group effect was tested by a Wald F-test, and the three planned contrasts were tested with emmeans using Holm adjustment within each endpoint. Kruskal-Wallis and Wilcoxon tests on the same three contrasts were run as a distribution-free sensitivity check.

Two additional models were used alongside this framework. Vein LDH was adjusted for pre-NMP baseline as a covariate (ANCOVA), since baseline varied widely across kidneys and predicted subsequent rise; its omnibus test used a likelihood-ratio comparison against a baseline-only model, since a standard F-test is not well-defined for a heteroscedastic ANCOVA. Time courses were also modeled by linear mixed-effects models (group × time, kidney random intercept) to characterize the overall trajectory, with planned contrasts evaluated at each outcome’s steady-state or 120-min summary value rather than from the trajectory model itself.

Histologic injury and Hosgood Quality Assessment scores are ordinal and sparsely distributed and were therefore analyzed by Kruskal–Wallis and Holm-adjusted Wilcoxon rank-sum tests for the same three contrasts, without a parametric model. For consistency with other outcomes, these scores are summarized as mean ± SD, and reported P values are Holm-adjusted.

The non-survival allotransplant cohort (fresh control, 24-h SCS and VNW, n = 3 per group; native contralateral kidneys pooled across groups, n = 9) was analyzed primarily descriptively, with exploratory hypothesis tests. For the 120-min endpoints and the steady-state urine production rate (60–120 min), VNW was compared with fresh control and with 24-h SCS using two-sided Welch’s t-tests, Holm-adjusted for the two comparisons. With n = 3 per group, the normality assumption could not be verified and the study was not powered for between-group inference. Native kidneys are shown as a reference and were not tested, because a native-versus-transplanted comparison requires a paired, within-animal analysis. Native urine production rate was derived from interval changes in cumulative urine volume.

## Supplementary Information

Supplementary Information is available for this paper.

### Correspondence

Correspondence and requests for materials should be addressed to Erik B. Finger or John C. Bischof.

## Acknowledgments

We thank John Carney and members of the University of Minnesota Experimental Surgical Services (ESS) and Advanced Preclinical Imaging Center (APIC) for assistance with organ recovery and transplantation; Matthew Powell-Palm and Markus Selzner for advice on the porcine transplant model; and Cameron Scheithauer, Seoha (Esther) Yoo, and Li Li for technical assistance. AI-assisted tools (Gemini, Google; ChatGPT, OpenAI; and Claude, Anthropic) were used during manuscript preparation for language editing, review, and formatting statistical outputs. All AI-assisted content was reviewed and verified by the authors, who take full responsibility for the final manuscript. The NMP perfusion circuit diagram was created using ChatGPT. This work was supported by National Institutes of Health grants DK131209 (E.B.F., J.C.B.), DK132211 (E.B.F., J.C.B.), DK117425 (E.B.F., J.C.B.), AI186974 (E.B.F., J.C.B.), DK120303 (E.B.F., J.C.B.), and HL185895 (Z.H.); National Science Foundation grant EEC 1941543 (E.B.F., J.C.B.); by the SAWIAGOS project; and by a gift from the Biostasis Research Institute, funded in part by contributions from LifeGift, Nevada Donor Network, Lifesource, Donor Network West, and Lifebanc organ procurement organizations.

## Author contributions

Z.H. and B.E.N. contributed equally to this work. J.C.B. and E.B.F. jointly supervised the work and contributed equally as senior authors. Conceptualization: M.L.E., E.B.F. and J.C.B.; methodology: M.L.E., E.B.F. and J.C.B.; investigation: all authors; visualization: Z.H., B.E.N., M.L.E., E.B.F. and J.C.B.; project administration: M.L.E., E.B.F. and J.C.B.; supervision: M.L.E., E.B.F. and J.C.B.; funding acquisition: E.B.F. and J.C.B.; writing, original draft: Z.H., B.E.N., M.L.E., E.B.F. and J.C.B.; writing, review and editing: all authors. All authors reviewed and approved the final manuscript.

## Data and materials availability

Raw data files are available on request.

## Extended Data Figures

**Extended Data Fig. 1.**
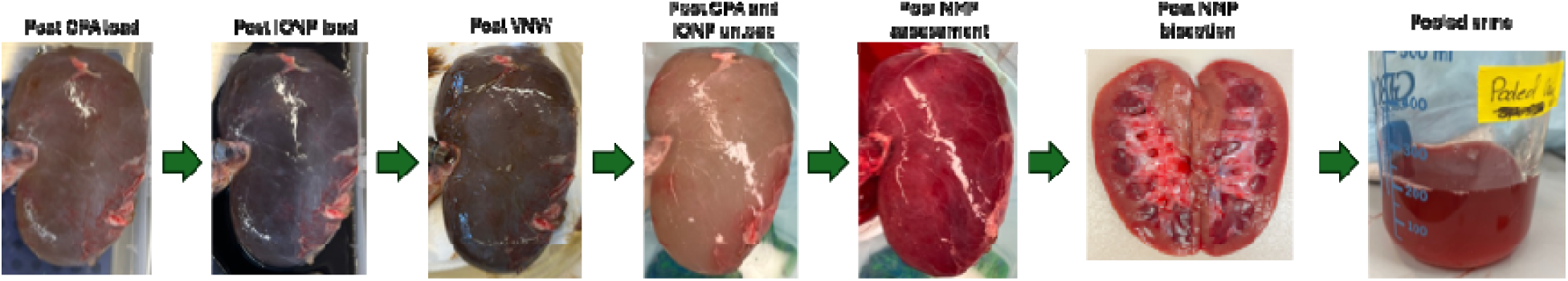
Gross appearance of a kidney throughout vitrification and nanowarming. Photographic sequence of a representative porcine kidney during CPA loading, vitrification, nanowarming and CPA unloading, followed by gross assessment after NMP, kidney bisection and urine produced during NMP. CPA, cryoprotective agent; sIONP, silica-coated iron oxide nanoparticle; VNW, vitrification and nanowarming; NMP, normothermic machine perfusion.

**Extended Data Fig. 2.**
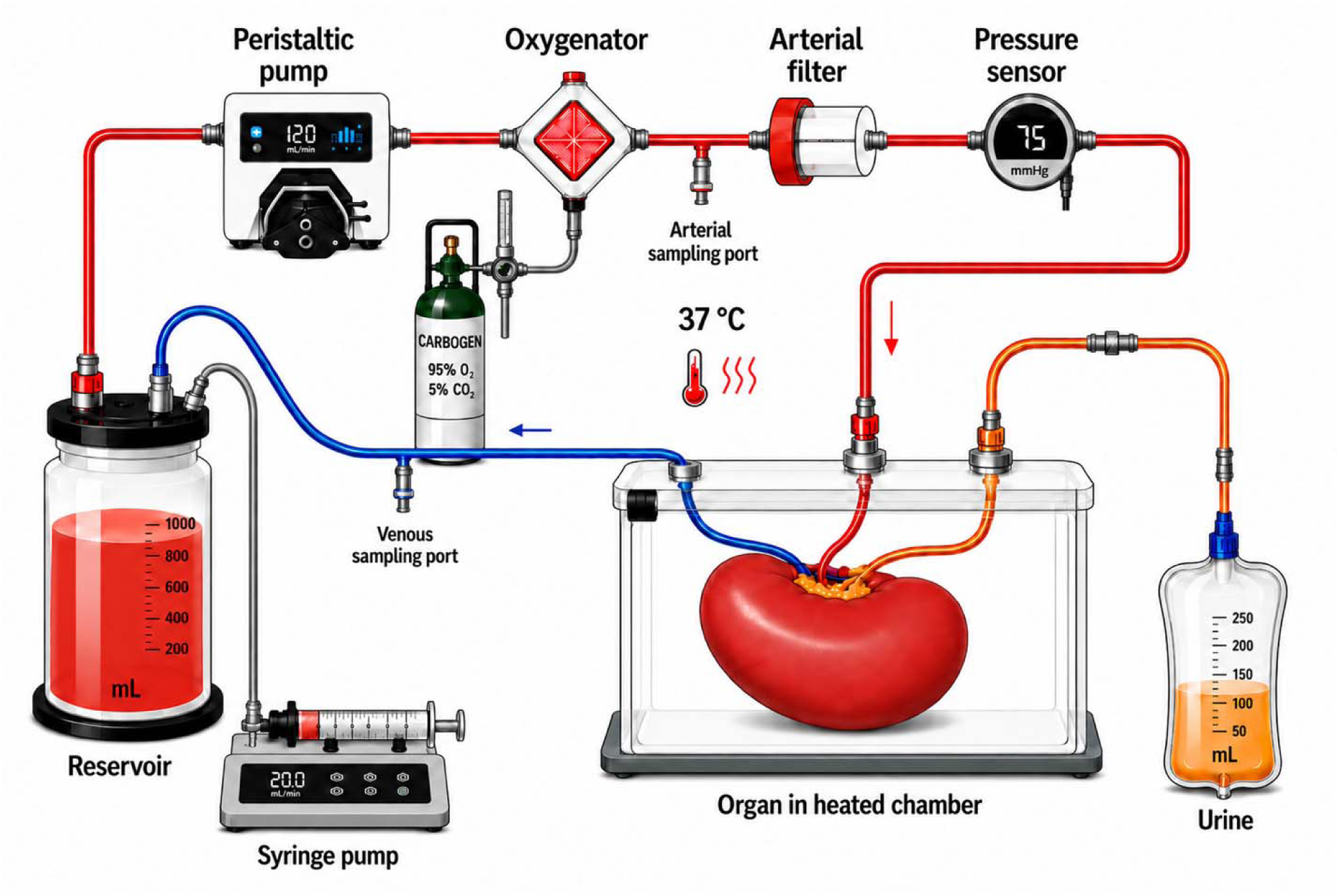
Normothermic machine perfusion (NMP) circuit for porcine kidney assessment. Schematic of the recycling NMP circuit used to assess recovered kidneys. Oxygenated autologous blood perfusate was circulated from the reservoir through a membrane oxygenator supplied with carbogen (95% O□/5% CO□), an arterial filter and an inline pressure sensor before entering the renal artery. Kidneys were maintained at 37 °C in a temperature-controlled chamber, and venous effluent was returned to the reservoir for recirculation. Arterial and venous sampling ports enabled serial assessment of metabolic and injury biomarkers, and a syringe pump was used to administer medications and supplements. Urine was collected separately for quantitative measurement and analysis.

**Extended Data Fig. 3.**
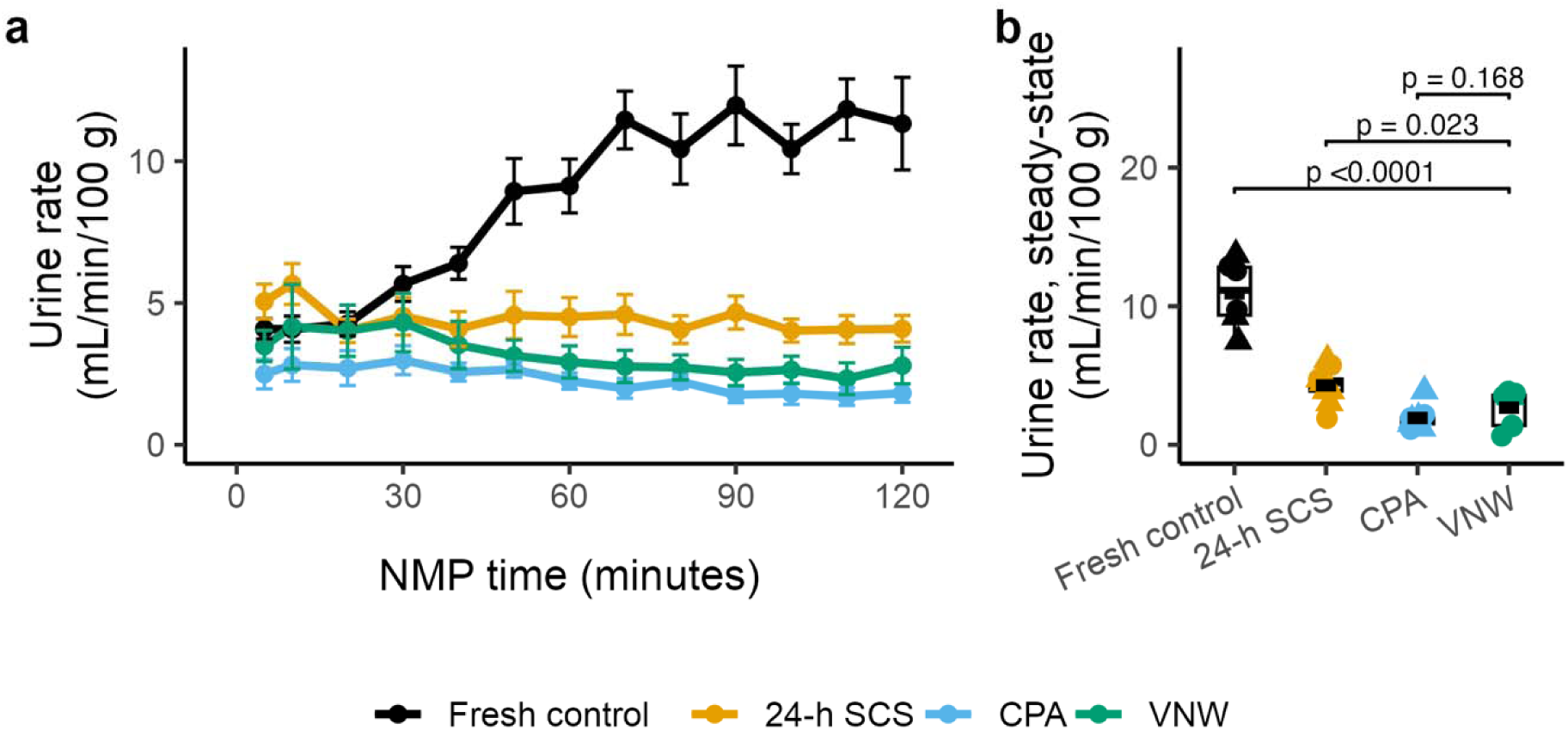
Urine production rate after vitrification and nanowarming. Urine production rate after vitrification and nanowarming (VNW) remained below that of fresh and 24-h static cold storage (SCS) kidneys and was not significantly different from cryoprotective agent (CPA)-only kidneys. CPA-only kidneys underwent CPA loading and unloading without vitrification, cryogenic storage or nanowarming. Biologically independent kidneys per group: fresh control, n = 6; 24-h SCS, n = 9; CPA-only, n = 9; VNW, n = 8 (both panels). **a,** Urine production rate over time; points and lines indicate group means and error bars indicate ± s.e.m. **b,** Steady-state urine production rate, calculated as the mean from 60–120 min; box-and-jitter plot conventions are as in Fig. 3. Brackets show prespecified comparisons of VNW with each other group. P values were calculated using a generalized least-squares (GLS) model allowing group-specific variance, followed by estimated marginal means with two-sided tests and Holm adjustment for the three comparisons; exact P values are shown above the brackets. Urine production rate was a secondary endpoint; cumulative urine production at 120 min (Fig. 3g) was the primary excretory endpoint.

**Extended Data Fig. 4.**
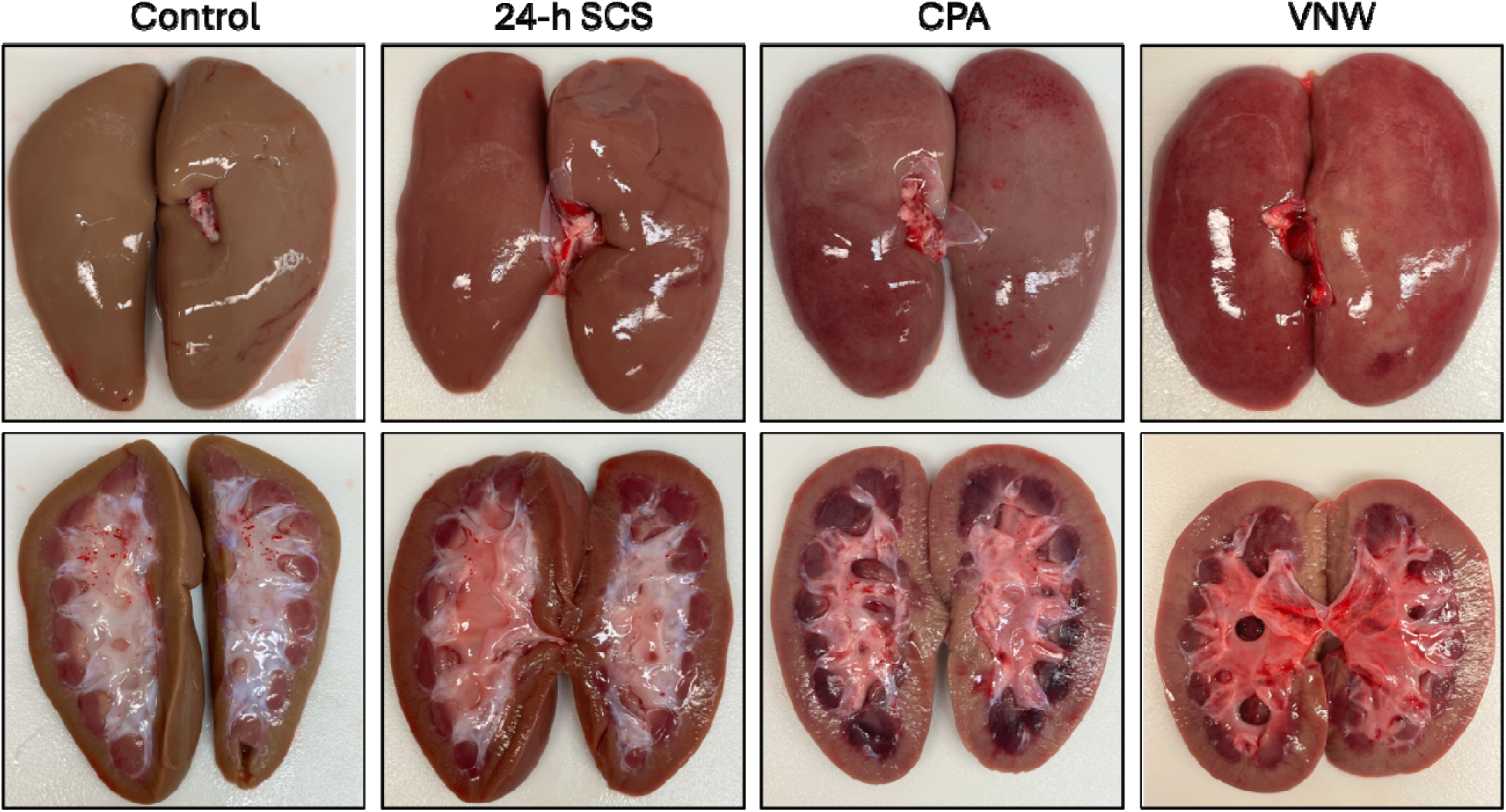
Gross appearance of kidneys following normothermic machine perfusion. Representative gross images of kidneys after 2 h of normothermic machine perfusion (NMP). Following NMP, the renal capsule was removed and each kidney was bisected in the coronal plane. Surface and cut-surface views are shown for fresh control, 24-h static cold storage (SCS), cryoprotective agent (CPA)-only and vitrification and nanowarming (VNW) groups.

**Extended Data Fig. 5.**
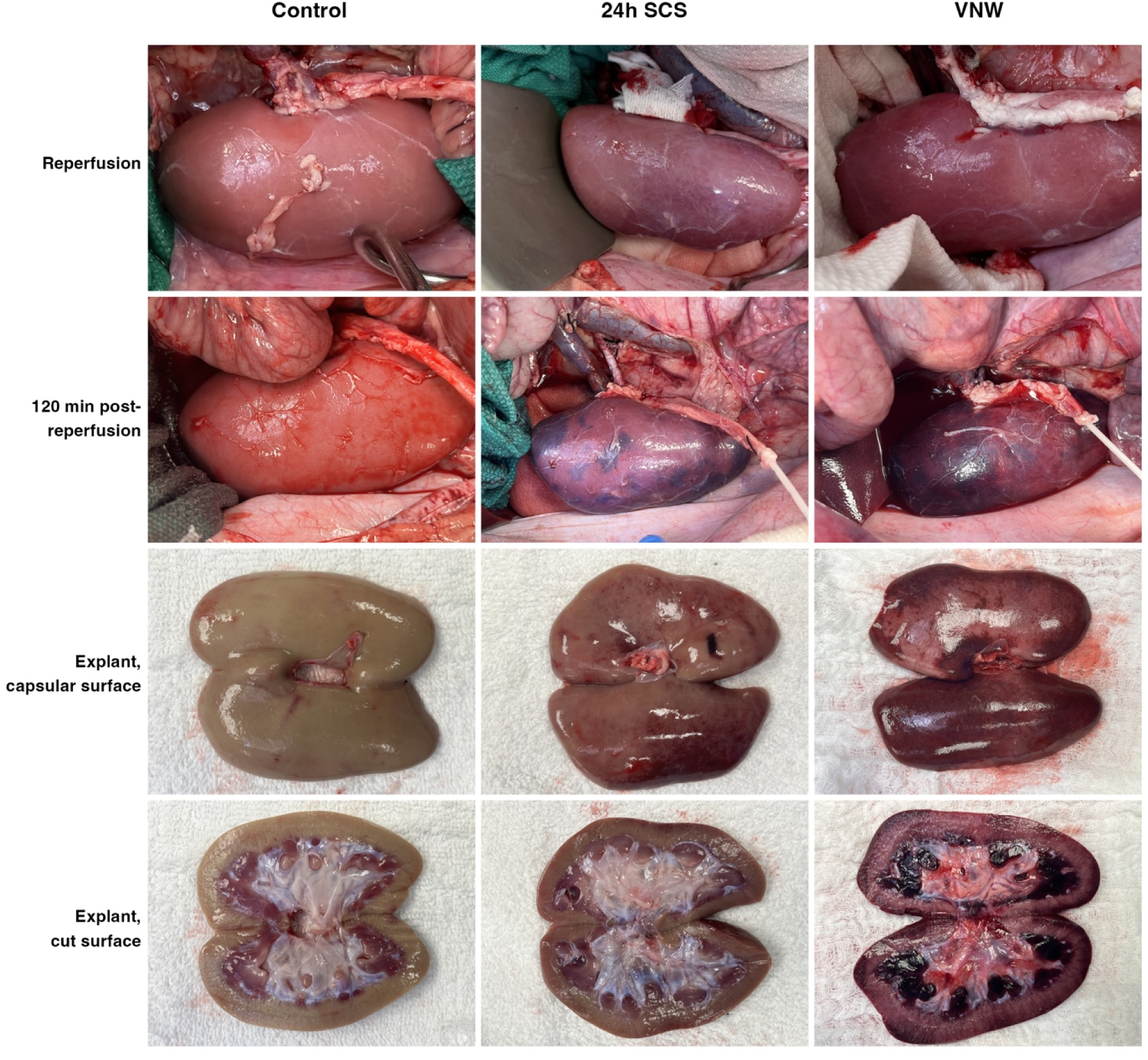
Gross appearance of transplanted kidneys after reperfusion. Representative images from the acute non-survival allotransplant cohort. Columns show fresh control, 24-h static cold storage (SCS) and vitrification and nanowarming (VNW) kidneys. Rows show the kidney immediately after reperfusion (intraoperative, in situ), at 120 min after reperfusion (intraoperative, in situ, end of study), after explantation (capsular surface, ex vivo) and after bisection (cut surface, ex vivo). One representative kidney from each group is shown at each stage. Images are illustrative and were not quantitatively assessed. The cohort comprised biologically independent kidneys from fresh control (n = 3), 24-h SCS (n = 3) and VNW (n = 3) groups.

## Extended Data Tables

**Extended Data Table 1.**
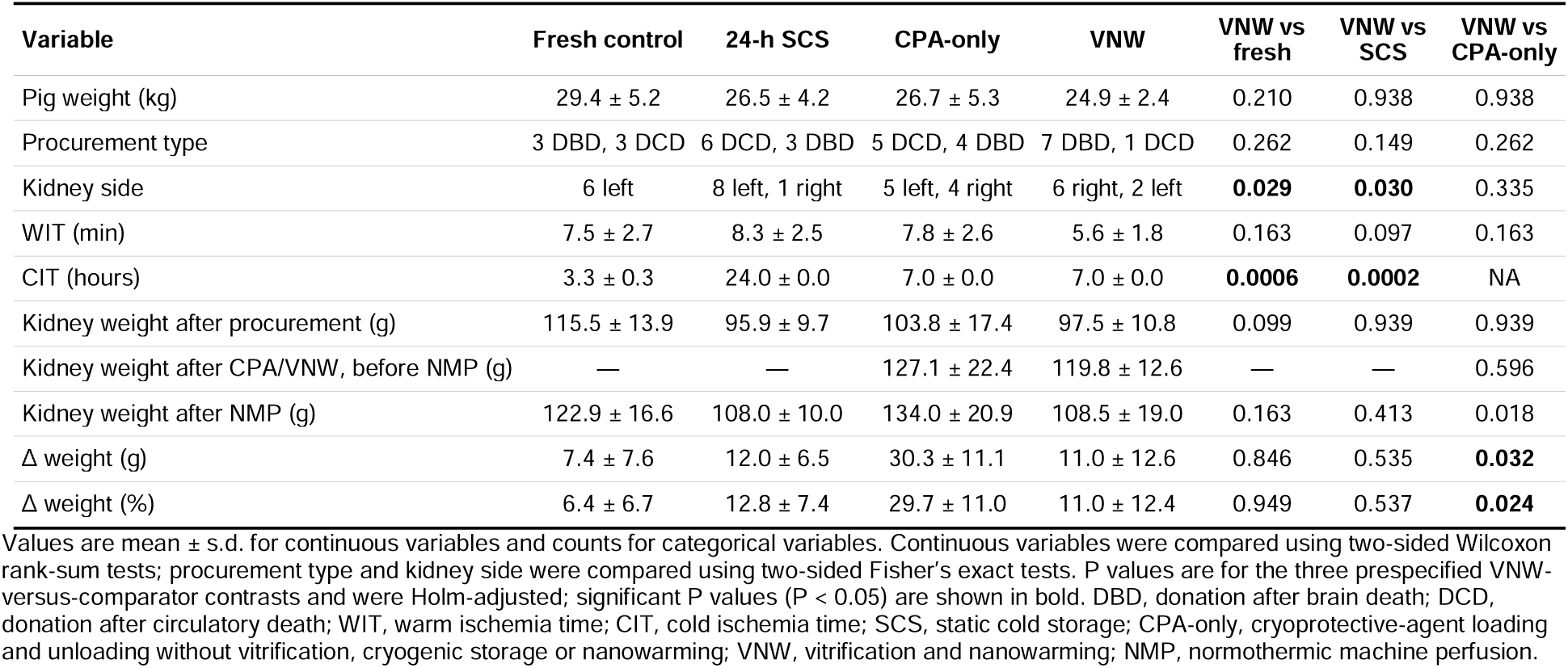
Demographic and procurement characteristics.

| Variable | Fresh control | 24-h SCS | CPA-only | VNW | VNW vs fresh | VNW vs SCS | VNW vs CPA-only |
| --- | --- | --- | --- | --- | --- | --- | --- |
| Pig weight (kg) | 29.4 ± 5.2 | 26.5 ± 4.2 | 26.7 ± 5.3 | 24.9 ± 2.4 | 0.210 | 0.938 | 0.938 |
| Procurement type | 3 DBD, 3 DCD | 6 DCD, 3 DBD | 5 DCD, 4 DBD | 7 DBD, 1 DCD | 0.262 | 0.149 | 0.262 |
| Kidney side | 6 left | 8 left, 1 right | 5 left, 4 right | 6 right, 2 left | **0.029** | **0.030** | 0.335 |
| WIT (min) | 7.5 ± 2.7 | 8.3 ± 2.5 | 7.8 ± 2.6 | 5.6 ± 1.8 | 0.163 | 0.097 | 0.163 |
| CIT (hours) | 3.3 ± 0.3 | 24.0 ± 0.0 | 7.0 ± 0.0 | 7.0 ± 0.0 | **0.0006** | **0.0002** | NA |
| Kidney weight after procurement (g) | 115.5 ± 13.9 | 95.9 ± 9.7 | 103.8 ± 17.4 | 97.5 ± 10.8 | 0.099 | 0.939 | 0.939 |
| Kidney weight after CPA/VNW, before NMP (g) | — | — | 127.1 ± 22.4 | 119.8 ± 12.6 | — | — | 0.596 |
| Kidney weight after NMP (g) | 122.9 ± 16.6 | 108.0 ± 10.0 | 134.0 ± 20.9 | 108.5 ± 19.0 | 0.163 | 0.413 | 0.018 |
| Δ weight (g) | 7.4 ± 7.6 | 12.0 ± 6.5 | 30.3 ± 11.1 | 11.0 ± 12.6 | 0.846 | 0.535 | **0.032** |
| Δ weight (%) | 6.4 ± 6.7 | 12.8 ± 7.4 | 29.7 ± 11.0 | 11.0 ± 12.4 | 0.949 | 0.537 | **0.024** |

Values are mean ± s.d. for continuous variables and counts for categorical variables. Continuous variables were compared using two-sided Wilcoxon rank-sum tests; procurement type and kidney side were compared using two-sided Fisher's exact tests. P values are for the three prespecified VNW-versus-comparator contrasts and were Holm-adjusted; significant P values (P < 0.05) are shown in bold. DBD, donation after brain death; DCD, donation after circulatory death; WIT, warm ischemia time; CIT, cold ischemia time; SCS, static cold storage; CPA-only, cryoprotective-agent loading and unloading without vitrification, cryogenic storage or nanowarming; VNW, vitrification and nanowarming; NMP, normothermic machine perfusion.

**Extended Data Table 2.** Functional, biochemical and histological outcomes.

| Outcome | Fresh control | 24-h SCS | CPA-only | VNW | P, VNW vs fresh | P, VNW vs 24-h SCS | P, VNW vs CPA-only |
| --- | --- | --- | --- | --- | --- | --- | --- |
| Flow, steady-state (ml/min/100 g) | 133.37 ± 35.55 | 135.70 ± 36.29 | 61.60 ± 32.78 | 81.67 ± 21.53 | **0.003** | **0.0005** | 0.132 |
| Resistance, steady-state (mmHg/ml/min/100 g) | 0.60 ± 0.20 | 0.60 ± 0.15 | 1.73 ± 0.97 | 1.04 ± 0.32 | **0.004** | **0.001** | **0.043** |
| Cumulative urine, 120 min (ml/100 g) | 1058.98 ± 232.29 | 525.68 ± 160.24 | 271.25 ± 85.10 | 375.59 ± 176.25 | **0.0006** | 0.177 | 0.177 |
| Urine rate, steady-state (ml/min/100 g) | 10.94 ± 2.47 | 4.29 ± 1.30 | 1.94 ± 0.81 | 2.68 ± 1.31 | **<0.0001** | **0.023** | 0.168 |
| Creatinine clearance, 30–60 min (ml/min/100 g) | 18.35 ± 12.50 | 12.56 ± 6.43 | 3.28 ± 1.80 | 4.16 ± 2.79 | **0.013** | **0.001** | 0.449 |
| O<sub>2</sub> consumption, steady-state (ml/min/100 g) | 0.47 ± 0.10 | 0.37 ± 0.04 | 0.26 ± 0.14 | 0.30 ± 0.08 | **0.001** | 0.069 | 0.454 |
| Venous pH, 120 min | 7.43 ± 0.02 | 7.42 ± 0.04 | 7.34 ± 0.04 | 7.34 ± 0.04 | **<0.0001** | **<0.0001** | 0.812 |
| Venous lactate, 120 min (mmol/l) | 0.62 ± 0.47 | 1.67 ± 0.81 | 1.53 ± 0.63 | 0.90 ± 0.39 | 0.231 | **0.034** | **0.034** |
| ATP, cortex, 120 min (nmol/mg protein) | 4.46 ± 0.60 | 4.40 ± 1.69 | 4.86 ± 2.35 | 4.96 ± 2.34 | 1.000 | 1.000 | 1.000 |
| ATP, medulla, 120 min (nmol/mg protein) | 25.91 ± 13.56 | 11.35 ± 9.44 | 8.08 ± 4.02 | 8.94 ± 4.09 | **0.044** | 1.000 | 1.000 |
| Urine NGAL, 120 min (ng/ml) | 6.51 ± 4.29 | 9.34 ± 8.73 | 138.35 ± 212.19 | 37.25 ± 10.84 | **<0.0001** | **<0.0001** | 0.154 |
| Venous LDH, 120 min (mU/ml)* | 1957.50 ± 744.00 | 1165.45 ± 611.98 | 1546.99 ± 643.23 | 1454.10 ± 316.43 | 1.000 | **0.008** | 1.000 |
| Histological injury, total (0–60)† | 6.50 ± 2.17 | 8.22 ± 2.33 | 10.11 ± 2.76 | 11.62 ± 2.97 | **0.019** | 0.091 | 0.264 |
| Quality Assessment Score, total (1–5)† | 1.00 ± 0.00 | 1.00 ± 0.00 | 2.56 ± 0.53 | 1.88 ± 0.35 | **0.005** | **0.001** | **0.014** |

Values are mean ± s.d. Biologically independent kidneys: n = 6/9/9/8 (fresh control/24-h SCS/CPA-only/VNW), except ATP (n = 4/8/8/7). P values are for the three prespecified VNW-versus-comparator contrasts and were Holm-adjusted; significant P values (P < 0.05) are shown in bold. Unless otherwise noted, contrasts are two-sided comparisons of estimated marginal means from generalized least-squares models with group-specific variances. \*Venous LDH was analyzed by generalized least-squares analysis of covariance with the pre-NMP baseline value as covariate. †Histological injury (total score) and Hosgood Quality Assessment Score (QAS; surface assessment) were analyzed by Kruskal–Wallis test with Holm-adjusted two-sided Wilcoxon rank-sum tests. SCS, static cold storage; CPA-only, cryoprotective-agent loading and unloading without vitrification, cryogenic storage or nanowarming; VNW, vitrification and nanowarming; NMP, normothermic machine perfusion; ATP, adenosine triphosphate; NGAL, neutrophil gelatinase-associated lipocalin; LDH, lactate dehydrogenase; QAS, Quality Assessment Score.

## Supplementary Figures

**Supplementary Fig. 1.**
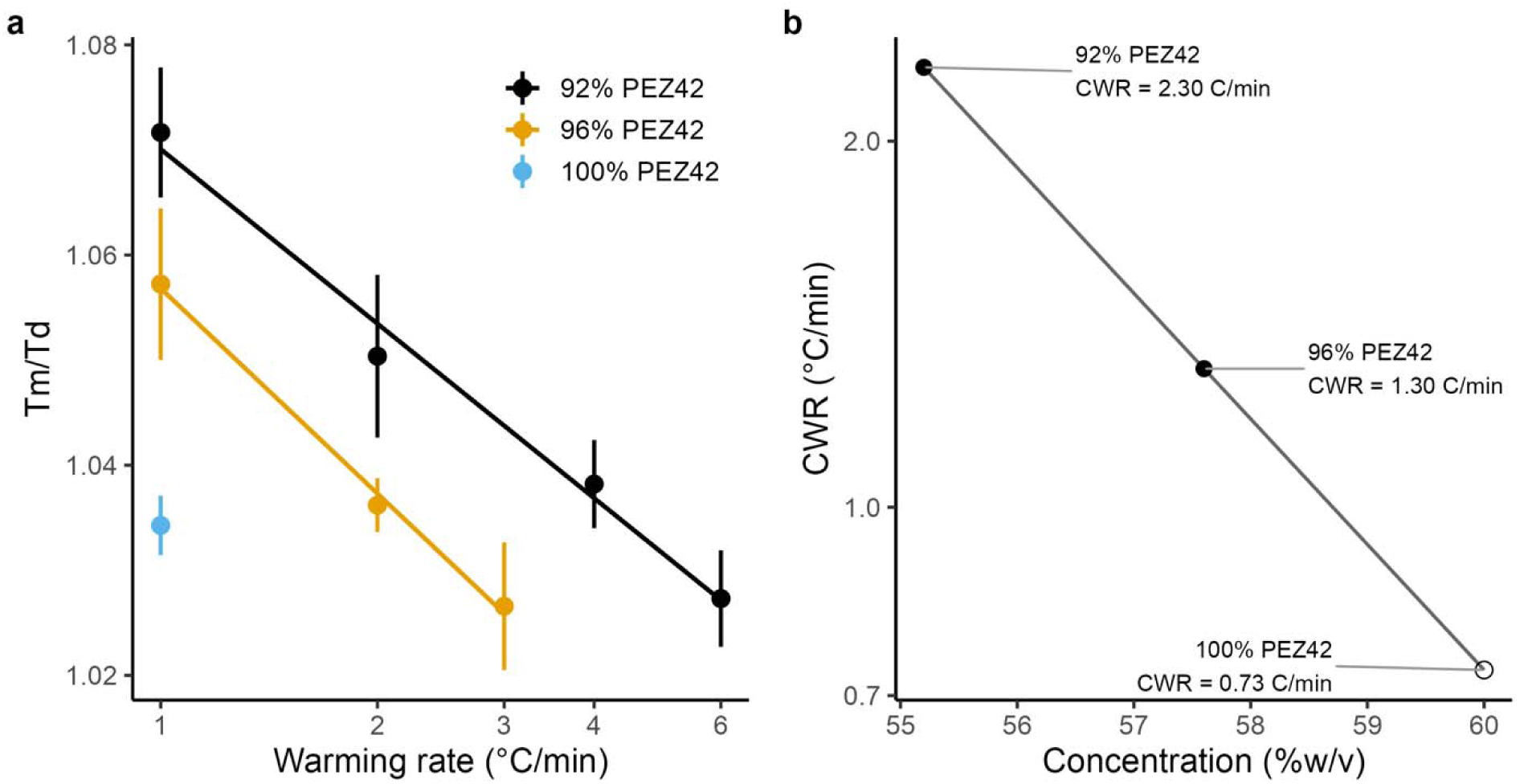
Critical warming rate (CWR) determination by differential scanning calorimetry. **a**, Ratio of melting temperature to devitrification temperature (Tm/Td) as a function of warming rate for PEZ42 at 92%, 96% and 100% of full-strength concentration (55.2, 57.6 and 60% w/v, respectively). Tm/Td approaches 1 as the devitrification interval decreases. Points show mean ± s.d. of 2–3 replicate DSC measurements at each warming rate. Lines show linear fits of Tm/Td versus log_10_(warming rate) for the 92% and 96% concentrations; the 100% concentration was evaluated at a single warming rate of 1 °C/min, which already gave ice crystallization below the 0.5% threshold and was therefore not fitted. **b**, Critical warming rate (CWR) as a function of PEZ42 concentration. Filled points indicate CWR values determined by regression of the DSC data using the 0.5% ice crystallization threshold; the open point indicates the calculated CWR for 100% PEZ42, for which ice crystallization is already below 0.5% at a warming rate of 1 °C/min. DSC, differential scanning calorimetry; CWR, critical warming rate.

**Supplementary Fig. 2.**
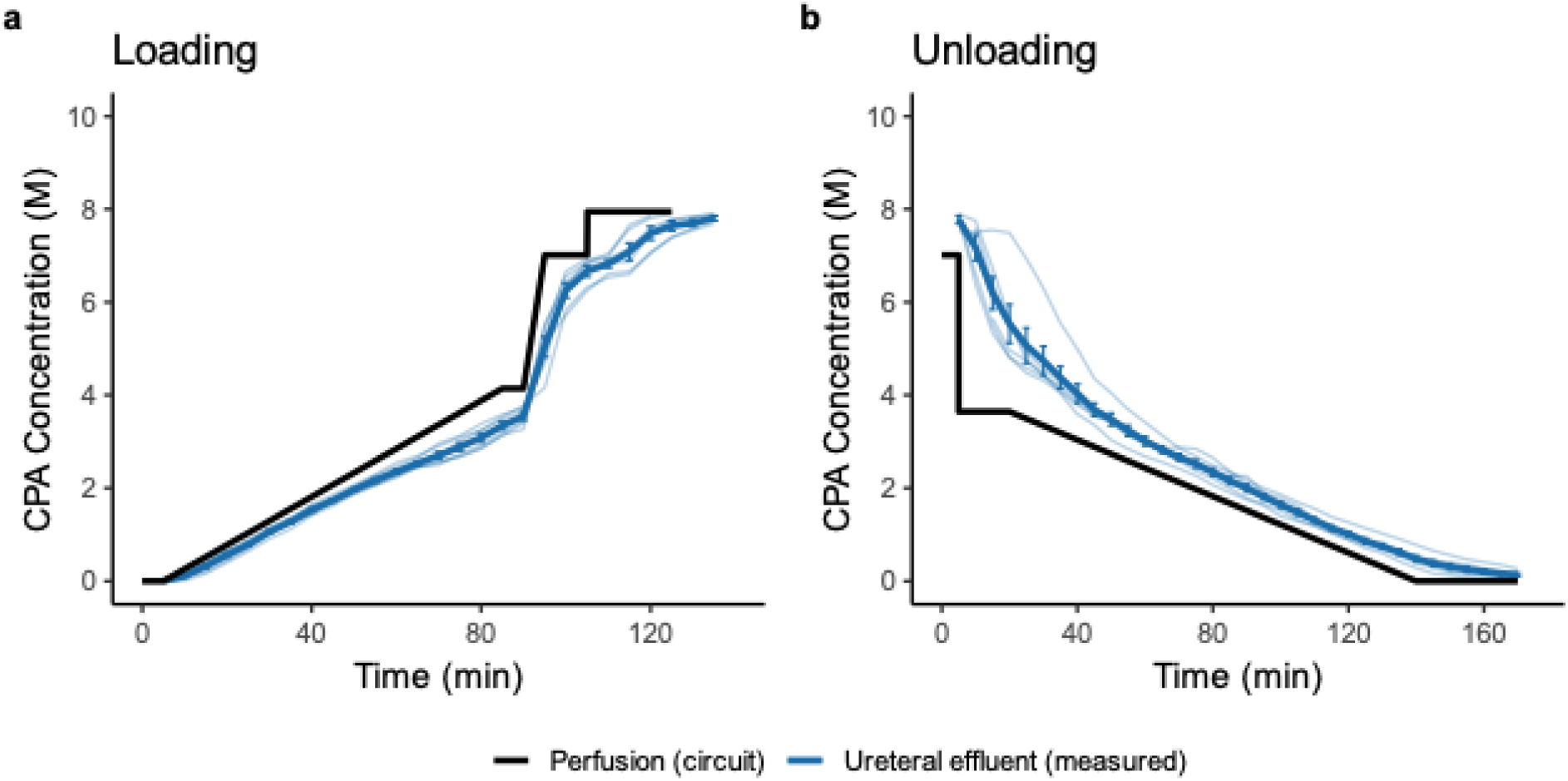
Ureteral effluent CPA concentration during loading and unloading. Perfusate cryoprotective agent (CPA) concentration specified by the loading and unloading protocol (black; as in Fig. 2a,c) is shown together with CPA concentration in ureteral effluent measured by refractometry (blue; thin lines, individual kidneys; thick line, mean; error bars, ± s.e.m.). **a,** CPA loading. Ureteral effluent CPA concentration increased with a delay relative to the perfusate concentration and was used as a surrogate for CPA equilibration within the renal medulla. To minimize exposure to high CPA concentrations, loading was terminated when ureteral effluent reached 97.5% of the full-strength CPA concentration. This feedback-guided endpoint reduced total loading duration from the initial fixed protocol of 135 min to 125 min and subsequently 120 min. **b,** CPA unloading. Ureteral effluent CPA concentration progressively decreased during CPA removal; unloading duration was fixed at 170 min for all six kidneys. n = 6 biologically independent kidneys.

**Supplementary Fig. 3.**
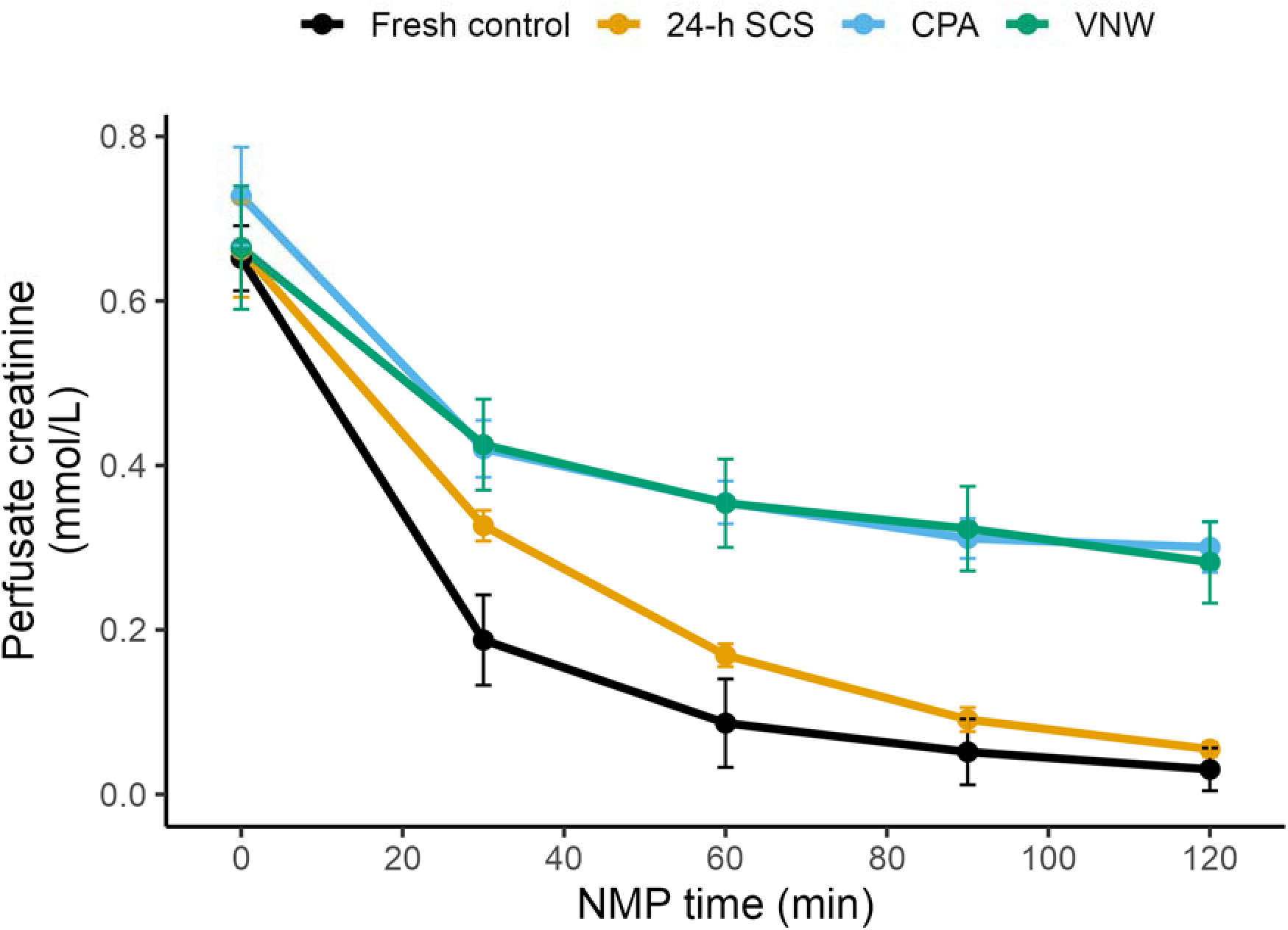
Perfusate creatinine decreases during normothermic machine perfusion. Perfusate creatinine concentration progressively decreased during normothermic machine perfusion (NMP), limiting interpretation of creatinine clearance at later time points. Biologically independent kidneys per group: fresh control, n = 6; 24-h static cold storage (SCS), n = 9; cryoprotective agent (CPA)-only, n = 9; vitrification and nanowarming (VNW), n = 8. Venous perfusate creatinine concentration is shown over time; points and lines indicate group means and error bars indicate ± s.e.m. Data are descriptive and no statistical comparisons are shown. The mean creatinine clearance from 30–60 min was therefore used as the primary clearance endpoint (Fig. 3h; see Methods).

**Supplementary Fig. 4.**
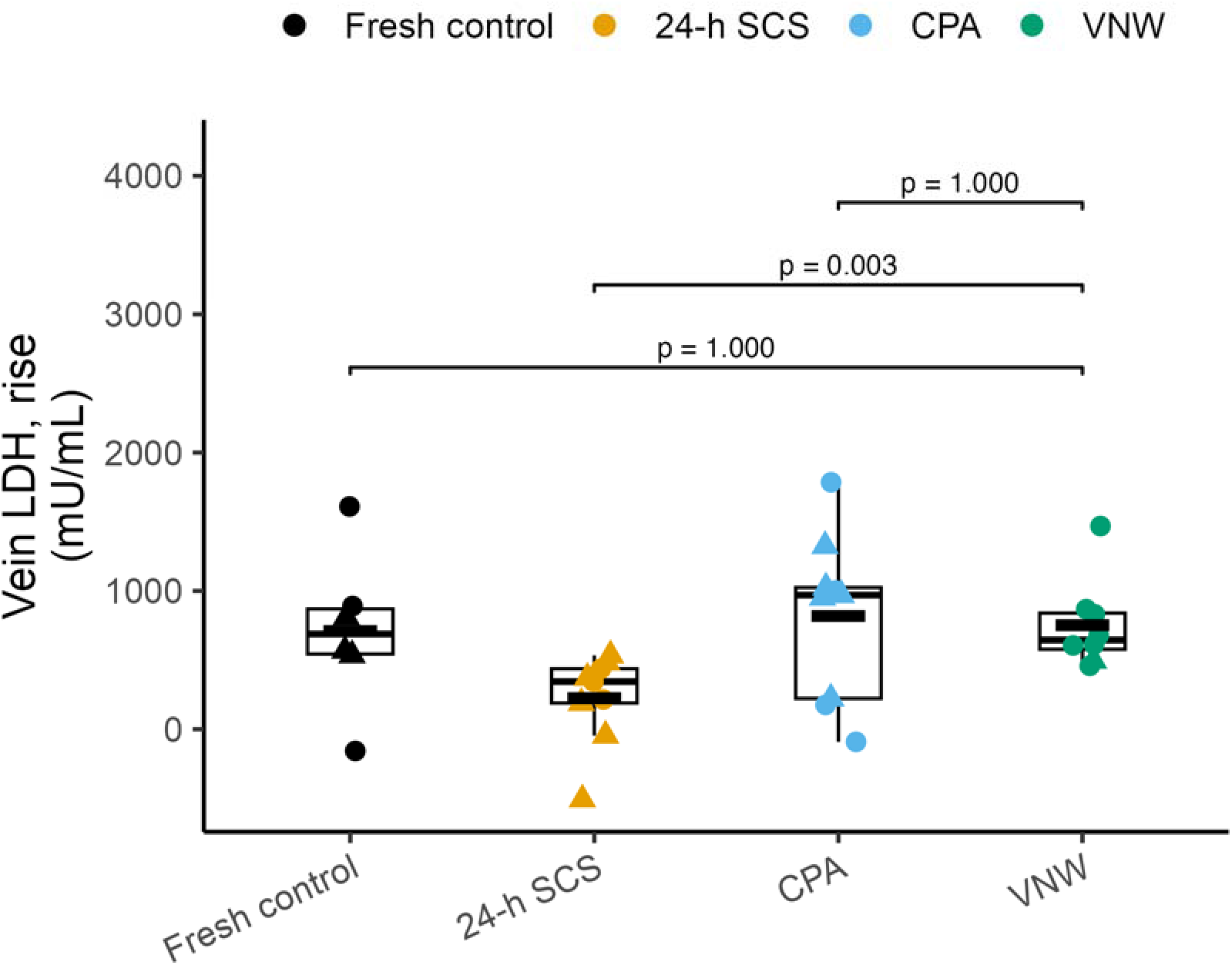
Absolute change in venous LDH during normothermic machine perfusion. Absolute change in venous lactate dehydrogenase (LDH) from baseline showed a pattern consistent with the primary baseline-adjusted LDH analysis. Cryoprotective agent (CPA)-only kidneys underwent CPA loading and unloading without vitrification, cryogenic storage or nanowarming. Biologically independent kidneys per group: fresh control, n = 6; 24-h static cold storage (SCS), n = 9; CPA-only, n = 9; vitrification and nanowarming (VNW), n = 8. Absolute change in venous LDH from the pre-NMP baseline to 120 min of normothermic machine perfusion (NMP) is shown. Box-and-jitter plot conventions are as in Fig. 3. Brackets show prespecified comparisons of VNW with each other group. P values were calculated using a generalized least-squares (GLS) model allowing group-specific variance, followed by estimated marginal means with two-sided tests and Holm adjustment for the three comparisons; exact P values are shown above the brackets. Absolute change in LDH was a secondary endpoint; baseline-adjusted LDH at 120 min (Fig. 4f) was the primary endpoint.

**Supplementary Fig. 5.**
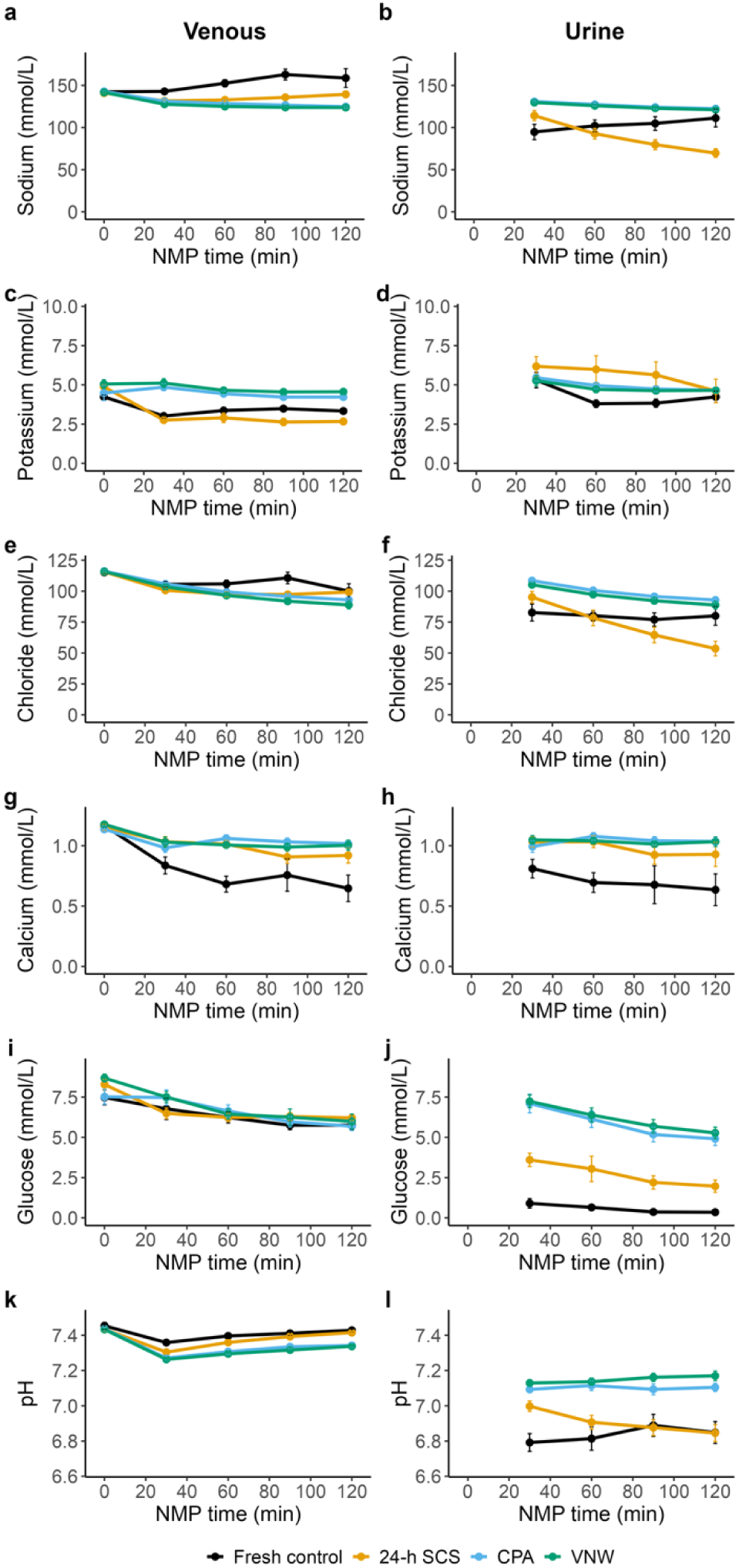
Perfusate and urine composition during normothermic machine perfusion. Perfusate and urine electrolyte, glucose and pH measurements were assessed across preservation groups during 120 min of normothermic machine perfusion (NMP). Cryoprotective agent (CPA)-only kidneys underwent CPA loading and unloading without vitrification, cryogenic storage or nanowarming. Biologically independent kidneys per group: fresh control, n = 6; 24-h static cold storage (SCS), n = 9; CPA-only, n = 9; vitrification and nanowarming (VNW), n = 8 (all panels). Sodium, potassium, chloride, calcium, glucose and pH are shown by row, with venous perfusate and urine measurements shown in the left and right columns, respectively. Measurements over time; points and lines indicate group means and error bars indicate ± s.e.m.

**Supplementary Fig. 6.**
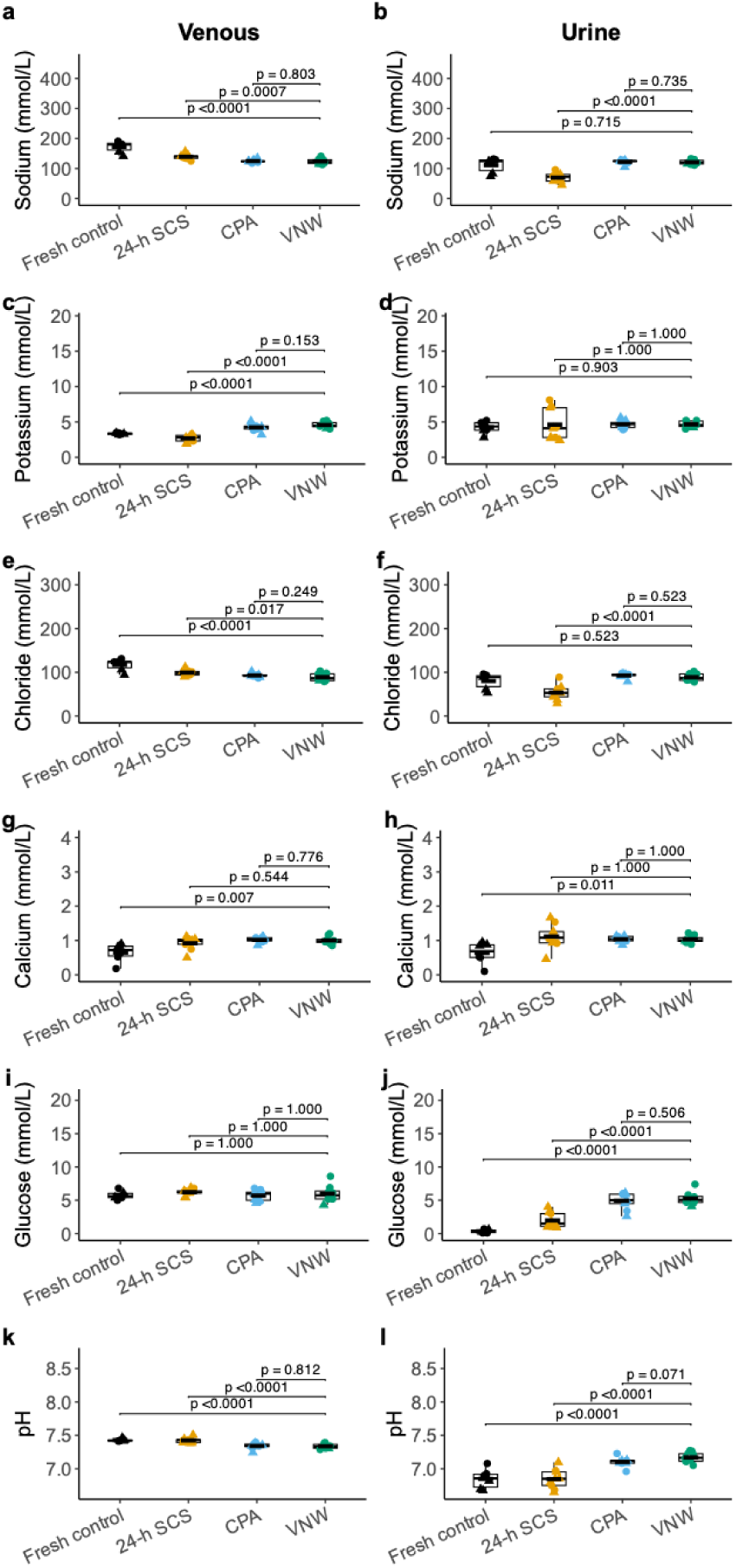
Endpoint perfusate and urine composition during normothermic machine perfusion. Perfusate and urine electrolyte, glucose and pH measurements were assessed across preservation groups during normothermic machine perfusion (NMP). Cryoprotective agent (CPA)-only kidneys underwent CPA loading and unloading without vitrification, cryogenic storage or nanowarming. Biologically independent kidneys per group: fresh control, n = 6; 24-h static cold storage (SCS), n = 9; CPA-only, n = 9; vitrification and nanowarming (VNW), n = 8 (all panels). Sodium, potassium, chloride, calcium, glucose and pH are shown by row, with venous perfusate and urine measurements shown in the left and right columns, respectively. Measurements at 120 min; box-and-jitter plot conventions are as in Fig. 3. Brackets show prespecified comparisons of VNW with each other group. P values were calculated using a generalized least-squares (GLS) model allowing group-specific variance, followed by estimated marginal means with two-sided tests and Holm adjustment for the three comparisons within each panel; exact P values are shown above the brackets.

**Supplementary Fig. 7.**
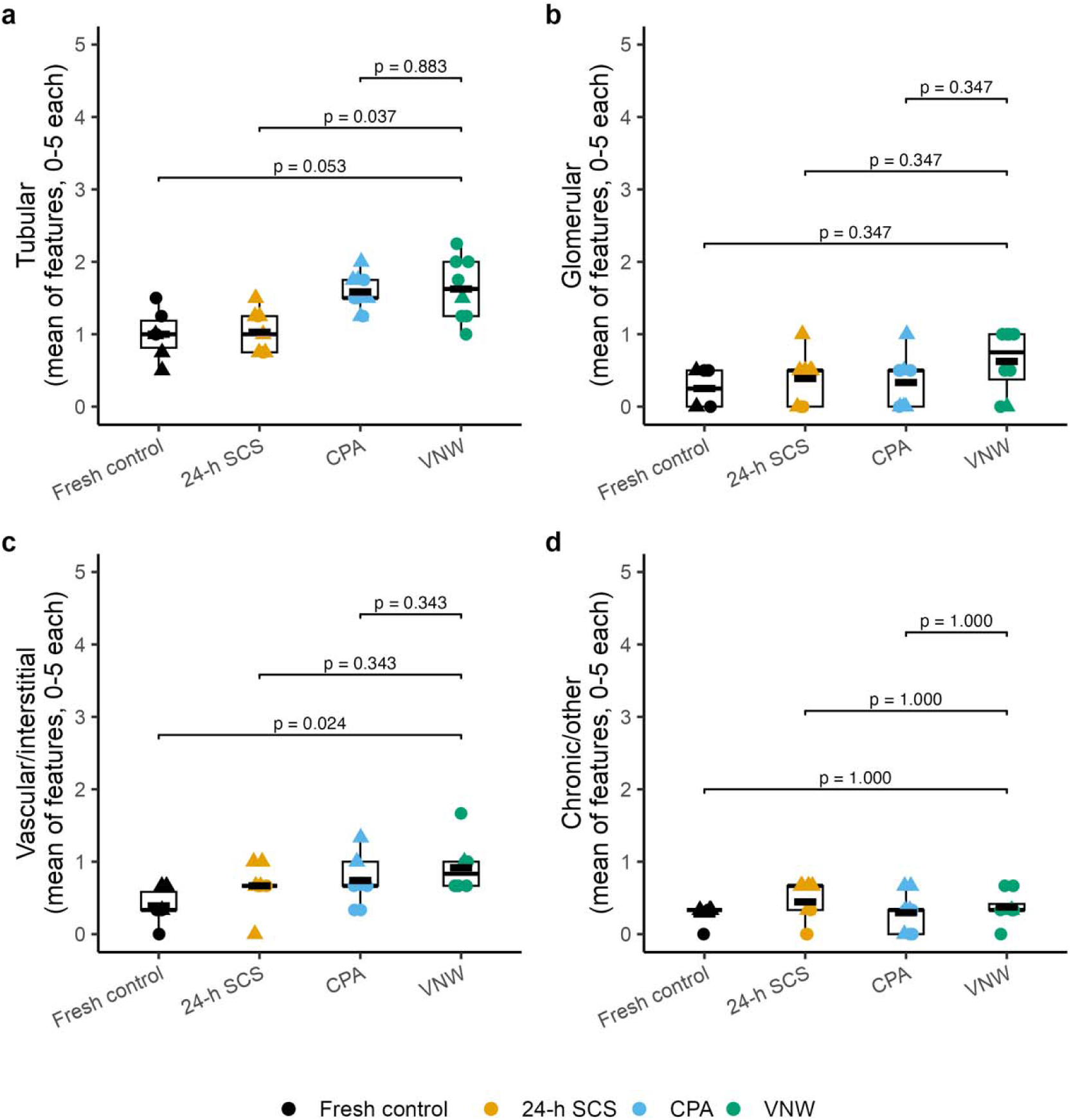
Histological injury is predominantly tubular after vitrification and nanowarming. Histological injury subscores show that injury after vitrification and nanowarming (VNW) is concentrated primarily in the tubular compartment. Cryoprotective agent (CPA)-only kidneys underwent CPA loading and unloading without vitrification, cryogenic storage or nanowarming. Biologically independent kidneys per group: fresh control, n = 6; 24-h static cold storage (SCS), n = 9; CPA-only, n = 9; VNW, n = 8 (all panels). Histological injury subscores represent the mean of component feature scores within each compartment (0–5 scale). Box-and-jitter plot conventions are as in Fig. 3. Brackets show prespecified comparisons of VNW with each other group. P values were calculated using two-sided Wilcoxon rank-sum tests with Holm adjustment for the three comparisons within each panel; exact P values are shown above the brackets. **a,** Tubular injury subscore. **b,** Glomerular injury subscore. **c,** Vascular/interstitial injury subscore. **d,** Chronic/other injury subscore.

**Supplementary Fig. 8.**
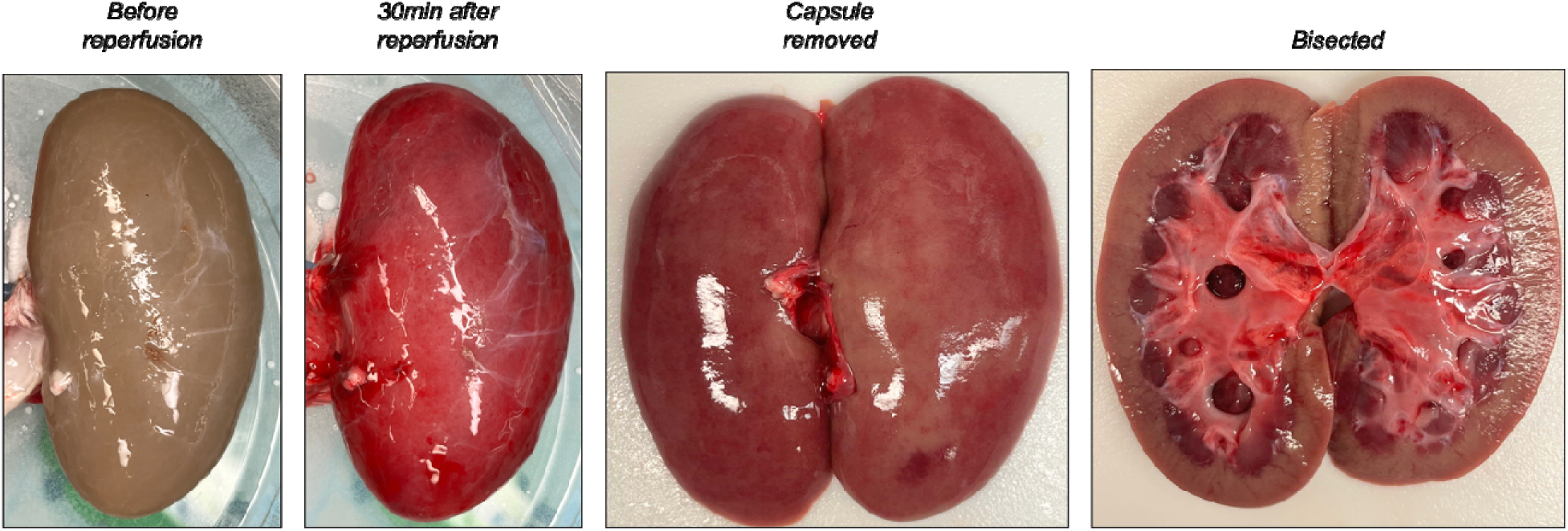
Surface erythema affects transplant Quality Assessment Score (QAS) but does not extend into renal cortex. Representative images of a vitrified and nanowarmed (VNW) kidney taken before normothermic machine perfusion, 30 min after initiation of reperfusion and at 120 min after capsule removal and bisection. The surface erythema affected Hosgood QAS determination but appeared superficial or subcapsular and did not extend into renal cortex. Reassessment based on bisected appearance improved QAS for CPA-only and VNW kidneys (Fig. 5d).

**Supplementary Fig. 9.**
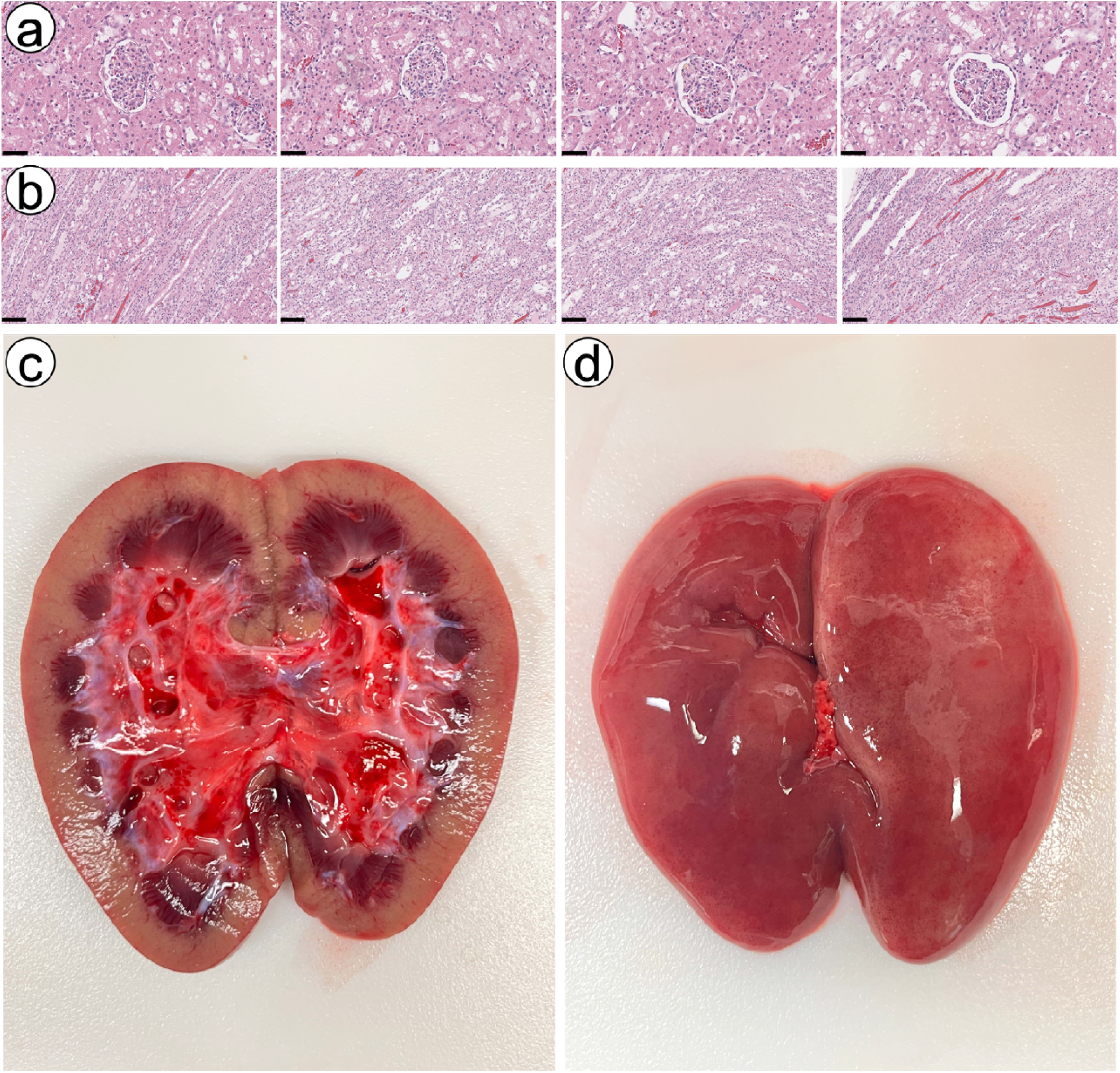
Histologic and gross appearance of a kidney after long-term cryopreservation. A single kidney was stored for 1 year (366 days) at −150 °C before nanowarming and normothermic machine perfusion (NMP). After 2 h of NMP, the renal capsule was removed and cortical and medullary samples were taken for histology. a, b, Renal cortex (a) and medulla (b) showing well-preserved histologic appearance. c, Gross appearance of the capsular surface and cut surface of the kidney, which also appeared well preserved. Scale bars, 50 µm (cortex) and 100 µm (medulla).

**Supplementary Fig. 10.**
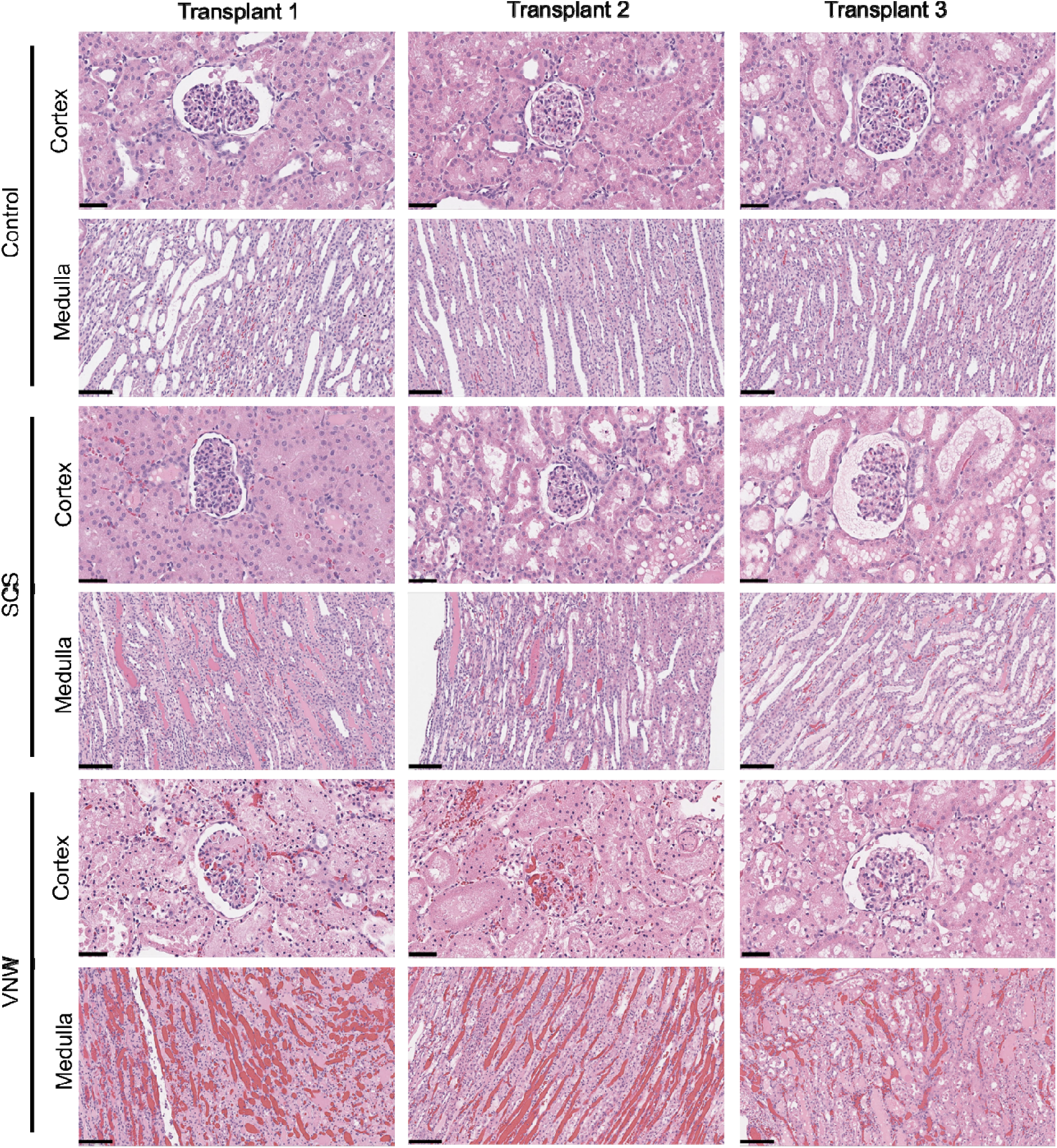
Histologic appearance of kidneys after acute allotransplantation. Representative cortical and medullary histology of fresh control, 24-h static cold storage (SCS) and vitrified and nanowarmed (VNW) kidneys 120 min after reperfusion in the acute non-survival allotransplant cohort. Images from three separate transplants are shown for each group. Scale bars, 50 µm (cortex) and 100 µm (medulla). Biologically independent kidneys: n = 3 per group.

**Supplementary Fig. 11.**
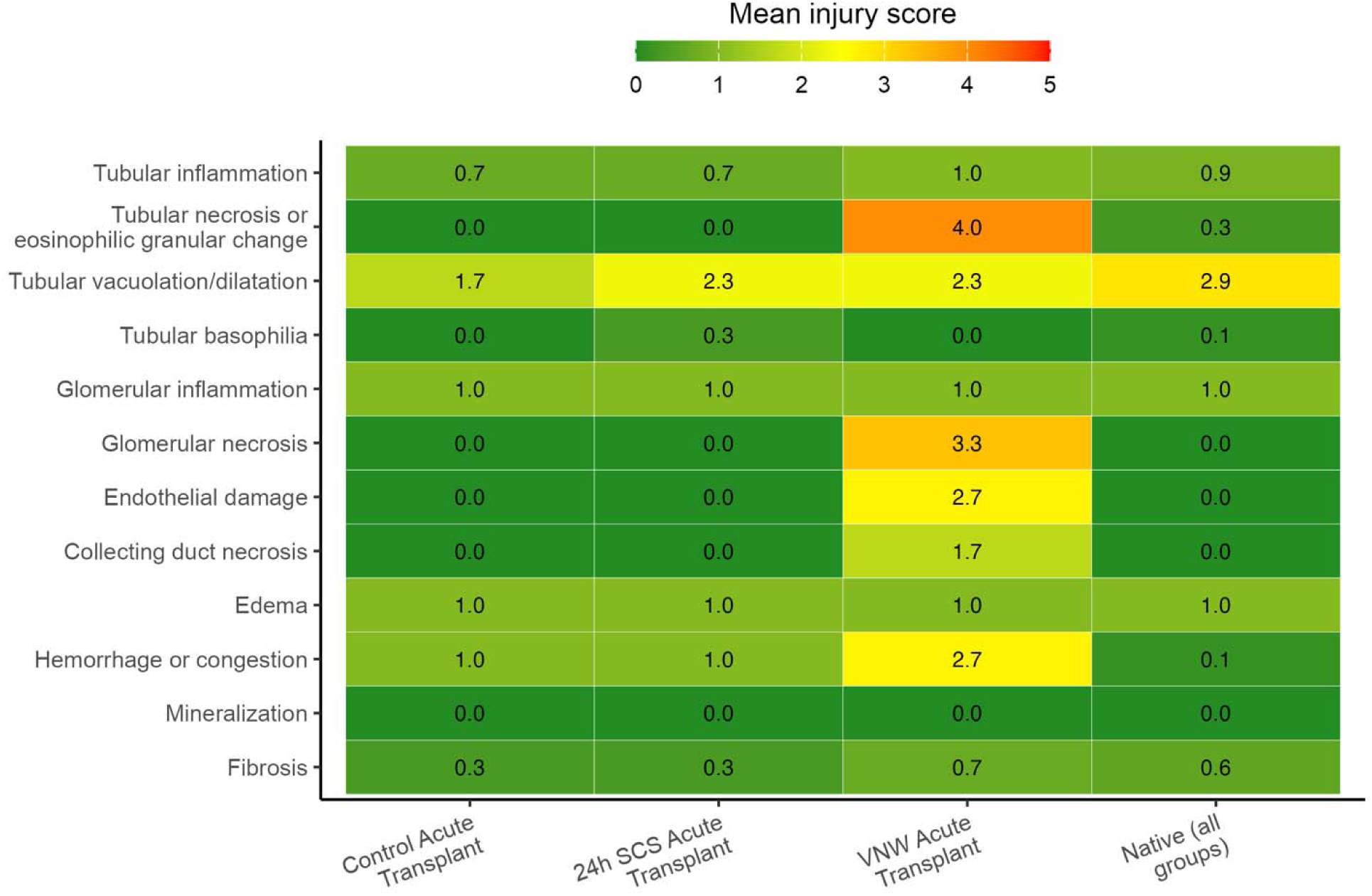
Histologic injury scores of kidneys after acute allotransplantation. **Heat map of mean histologic injury scores for the 12 scored features (rows) by group (columns) in the acute non-survival allotransplant cohort. Donor kidneys were** transplanted immediately after procurement (fresh control, n = 3), after 24-h static cold storage (SCS; n = 3) or after vitrification and nanowarming (VNW; n = 3); native contralateral kidneys from seven of the nine recipients (n = 7; two were not sampled) were pooled across all three groups and are shown as a reference. Each feature was scored on a 0–5 scale by a blinded pathologist; shading and the value printed in each cell indicate the group mean. Data are descriptive and no statistical comparisons are shown.

## Supplementary Tables

**Supplementary Table 1.**
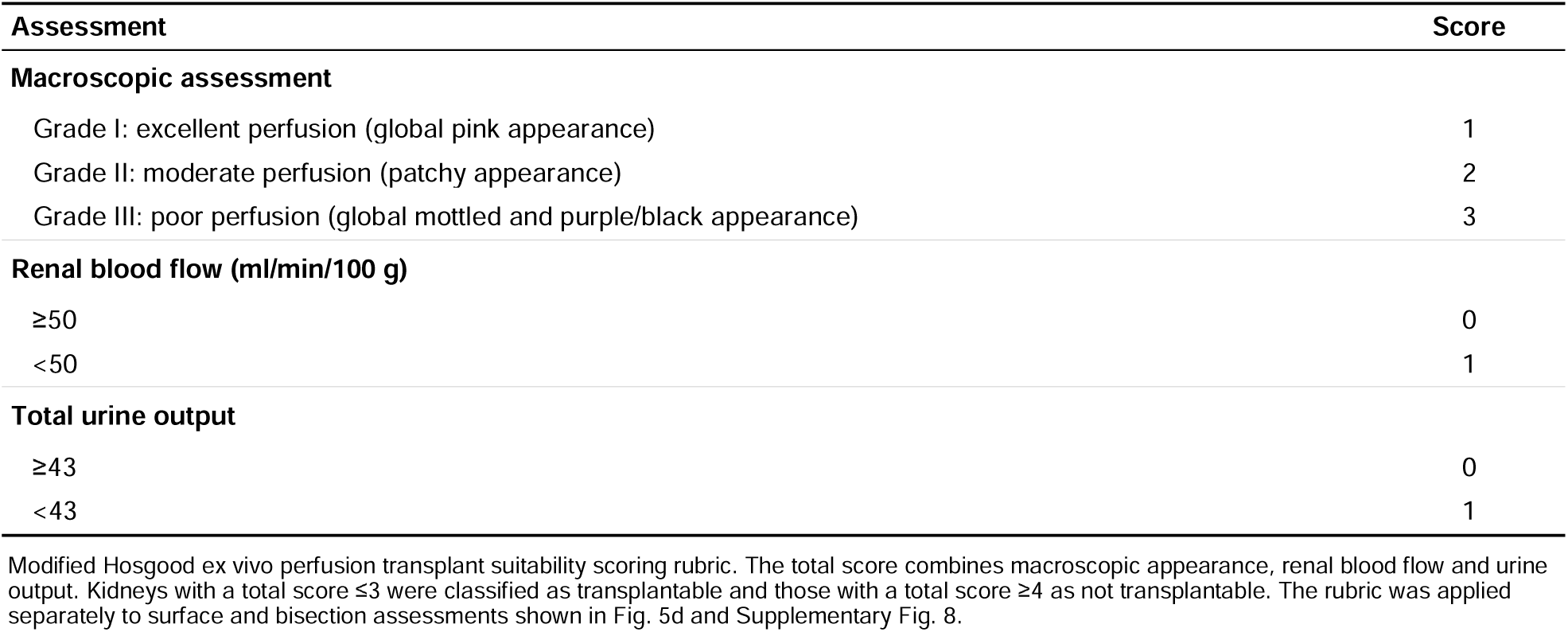
Hosgood transplant suitability score.

**Supplementary Table 2.** Histological injury scoring rubric.

| Severity | Proportion of tissue affected | Grade | Quantifiable finding |
| --- | --- | --- | --- |
| None | 0% | 0 | 0 |
| Marginal or minimal | 1–10% | 1 | 1–2 foci |
| Slight or few | 11–25% | 2 | 3–6 foci |
| Moderate or several | 26–50% | 3 | 7–12 foci |
| Marked or many | 51–75% | 4 | >12 foci |
| Severe | >75% | 5 | Diffuse |

## Supplementary Videos

**Supplementary Video 1 Rapid reperfusion and urine production of vitrified and nanowarmed (VNW) kidneys during normothermic machine perfusion (NMP).** Representative video of a kidney cryopreserved for 1 year showing rapid and homogeneous reperfusion and immediate urine production after initiation of perfusion at 37 °C during NMP.

**Supplementary Video 2 Rapid reperfusion and urine production of vitrified and nanowarmed (VNW) kidneys after allotransplantation. Representative video showing rapid and homogeneous** reperfusion and immediate urine production after initiation of reperfusion.

